# Involvement of nucleus accumbens D2-MSN projections to the ventral pallidum in anxious-like behavior

**DOI:** 10.1101/2022.05.23.493058

**Authors:** Raquel Correia, Ana Verónica Domingues, Bárbara Coimbra, Natacha Vieitas-Gaspar, Nuno Sousa, Luísa Pinto, Ana João Rodrigues, Carina Soares-Cunha

## Abstract

The nucleus accumbens (NAc) is a crucial brain region for emotionally-relevant behaviors. The NAc is mainly composed of medium spiny neurons (MSN) expressing either dopamine receptor D1 (D1-MSNs) or D2 (D2-MSNs). D1-MSNs project to the ventral tegmental area (VTA) and ventral pallidum (VP), while D2-MSNs project only to the VP. In this work, we selectively manipulated D1-MSN projections to the VP and VTA, and D2-MSN projections to the VP during classical anxiety behavioral paradigms in naïve mice.

We found that optogenetic activation of D1-MSN to VP or VTA did not trigger significant anxious-like behaviors. Interestingly, optical activation of D2-MSN-VP projections significantly increased anxious-like behavior in all of the tests performed. This phenotype was associated with a decrease in the activity of VP putative GABAergic neurons. Importantly, pre-treating D2-MSN-VP animals with the GABA modulator diazepam prevented the optically-triggered anxious-like behavior.

Overall, our results suggest that D2-MSN-VP projections contribute for the development of anxious-like behavior, through modulation of GABAergic activity in the VP.

## Introduction

Increasing evidence supports the existence of alterations in brain regions of the reward circuit in neuropsychiatric disorders such as depression and anxiety (Russo and Nestler, 2013). Importantly, both disorders are characterized by impaired responses to rewarding and aversive events (Hein et al., 2021; Satterthwaite et al., 2015), hinting for a central role of the reward circuit in the emergence of both depressive and anxiety symptoms (Russo and Nestler, 2013). One of the key regions of the reward circuit is the nucleus accumbens (NAc), which is anatomically positioned to integrate and convey limbic and motor information to drive behavior in response to emotionally relevant events (Morrison et al., 2017).

Increasing evidence shows the involvement of the NAc in depressive-like behaviors (Bewernick et al., 2012; Epstein et al., 2006; Francis et al., 2015; Lim et al., 2012; Salamone and Correa, 2012); however, the role of this brain region in anxious-like behavior is less explored. Still, recent studies suggest an important role of the NAc in the etiology and maintenance of aberrant avoidance behaviors in anxiety disorders. For example, in an avoidance task, the degree of activation and deactivation of the NAc was associated with individual levels of anxiety (Levita et al., 2012). In addition, a larger bilateral NAc volume is observed among adults with generalized anxiety disorder and is associated with higher levels of trait anxiety (Kühn et al., 2011), and NAc volume is a predictor of patient response to either cognitive-behavioral therapy or pharmacological treatment (Burkhouse et al., 2020). In addition, association between NAc taurine levels and trait anxiety have been observed (Berchio et al., 2019). In accordance, in animal models of anxiety, pharmacological and genetic modulation of NAc activity results in regulation of anxiety-like behavior (Feng et al., 2017; Heshmati et al., 2016; Shen et al., 2016). The abovementioned findings are not surprising, considering that the NAc is innervated by the amygdala (Li et al., 2018), a key region for anxiety (Li et al., 2018). In addition, NAc outputs to the VTA are key mediators of depressive and anxious-like phenotypes (Felix-Ortiz et al., 2016; Tye et al., 2011; Zweifel et al., 2011), with NAc-VTA circuit underlying emotional stress-induced anxiety-like behavior (Qi et al., 2022).

The NAc is mainly composed of medium spiny neurons expressing dopamine receptor D1 (D1-MSNs) or D2 (D2-MSNs). D1-MSNs directly project to the ventral tegmental area (VTA) (direct pathway), and the ventral pallidum (VP) is innervated by both D1- and D2-MSNs (indirect pathway) (Kupchik et al., 2015). The VP then projects to the VTA through GABAergic projecting neurons (Kupchik et al., 2015). While there is some consensus about the differential role of these two subpopulations in depressive-like behaviors, the contribution of D1- and D2-MSN neurons in anxiety traits is unknown.

Given this, in this work, we evaluated how D1-MSN projections to the VP or VTA, or D2-MSN projections to the VP contribute for the development of anxious-like behaviors in mice. We optogenetically activated these projections during classical behavioral paradigms to observe if one could attenuate or trigger an anxiety-like phenotype. We also characterized the electrophysiological response of downstream target regions, the VP and VTA, to D1- or D2-MSN optical activation. Our data show that activation of D2-MSN-VP outputs induces robust anxious-like behavior, and that this phenotype is reversed by treating animals with diazepam, a potent GABA modulator.

## Results

To explore if D1-MSN outputs to the VTA or VP, or D2-MSN projections to the VP could contribute for the development of anxious-like behaviors, we optogenetically activated (20Hz, 25ms light pulses) these projections during a battery of classical anxiety behavioral tests. To guarantee that we were able to compare performance within the same animal between optical stimulation (ON) epochs and no optical stimulation (OFF) epochs, we used previously published versions of the behavioral tasks that included alternating epochs of OFF-ON-OFF periods (Felix-Ortiz et al., 2016; Kim et al., 2013). When this option was not possible due to task design, the same behavioral test was performed twice for each animal (ON and OFF sessions, order counterbalanced).

### Optical activation of D1-MSN projections to the VP induces a very mild anxious-like behavior

To investigate if D1-MSN-VP projections are involved in the induction of anxious-like behavior, we unilaterally injected a cre-dependent construct containing channelrhodopsin (ChR2) in fusion with the enhanced yellow fluorescent protein (YFP) (AAV5-EF1A-DIO-hChR2(H134R)-eYFP), or a control YFP virus, in the NAc of D1-cre male mice (Figure 1A). We placed an optical fiber in the VP, to allow optical activation of D1-MSN terminals in the VP during behavior. Nearly 30% of the VP area presented YFP innervation (D1-MSN terminals) (Figure 1B-C).

**Figure 1.**
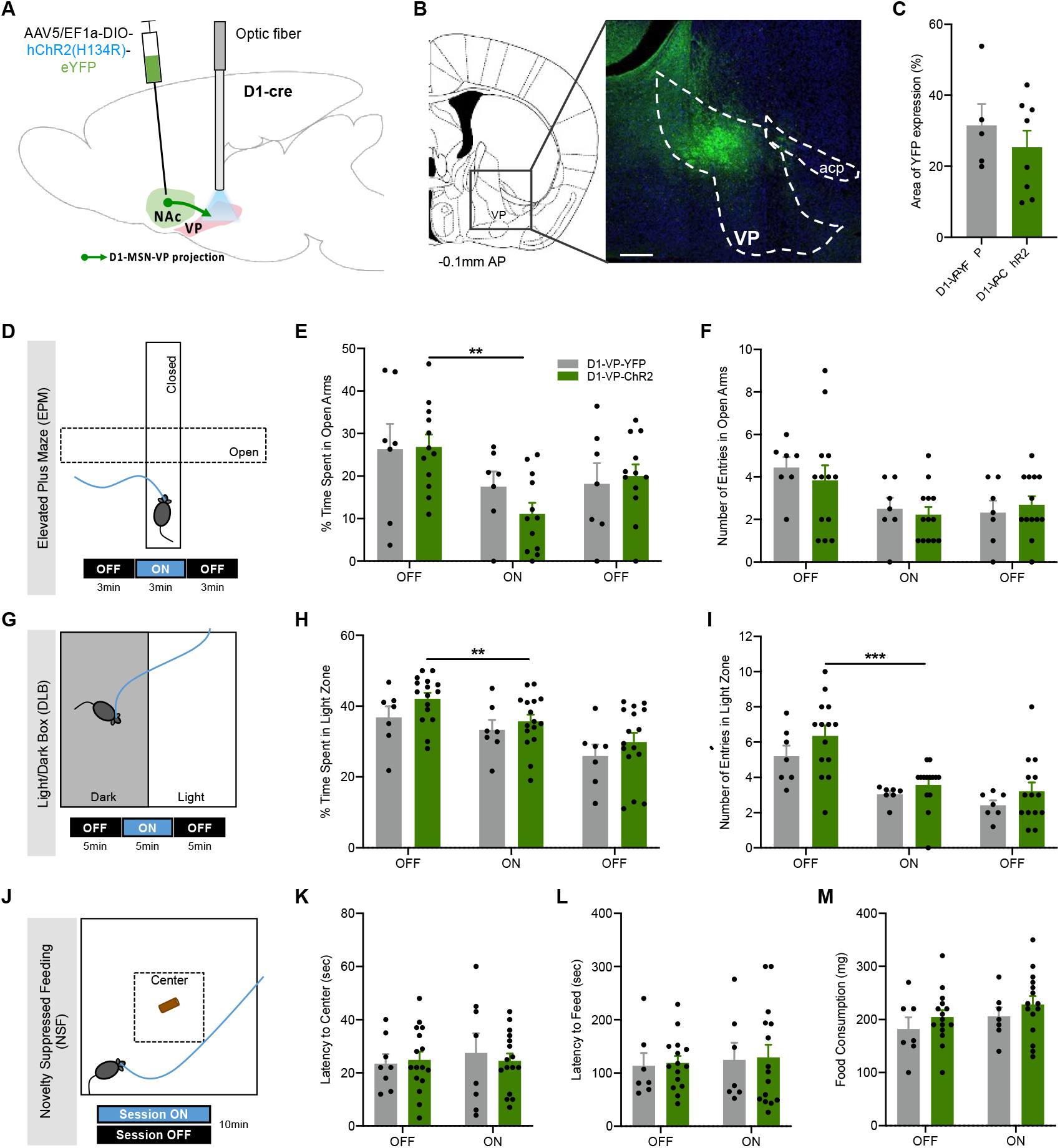
Optogenetic stimulation of D1-MSNs projecting to the VP in anxious-like behavior. **A** D1-cre mice were injected with either cre-dependent ChR2 or YFP. Blue light was delivered through an optical fiber implanted in the VP to allow terminal stimulation (473nm, 20Hz, 25ms pulses, 50% duty cycle). **B** Coronal brain slice showing expression of YFP using immunostaining. Numbers represent distance to bregma in millimeters; AP: anteroposterior; scale bar: 500 μm. **C** About 30% of VP region presented YFP expression, indicative of D1-MSN terminals. **D** The EPM was divided in 3-min epochs: OFF-ON-OFF. D1-MSN-VP optical stimulation induced slight decrease in the **E** time spent in the open arms, without altering the **F** number of entries in the open arms. **G** The LDB test was divided in 5-min epochs: OFF-ON-OFF. No differences were observed in the **H** time spent or **I** number of entries in the light zone of the arena. **J** Two sessions of NSF were performed: 1 session ON and 1 session OFF (counterbalanced); optical stimulation was performed throughout the entire session ON. D1-VP-ChR2 mice showed a similar **K** latency to reach the food pellet located in the center of the arena and **L** to feed in the ON and OFF sessions. **M** food consumption after the NSF was not altered. Data denote mean±SEM. *p≤0.05, **p≤0.01, ***p≤0.001.

D1-VP-ChR2 mice spent less time in the open arms of the EPM test during the ON period of the session in comparison with the OFF period (F_2,22_=8.8, *p*=0.0016; OFF vs ON p=0.002). However, the percentage of time spent in the open arms in the ON session was similar between ChR2 and YFP control mice (*p*=0.285) (Figure 1D-E; Figure 1 – figure supplement 1A). No significant differences were observed in the number of entries in the open arms of the maze (Figure 1F; Figure 1 – figure supplement 1B). In the LDB test, D1-VP-ChR2 mice spent a similar amount of time (F_1,15_=2.3, *p*=0.149) and entered in the light zone in similar levels to D1-VP-YFP control animals (F_1,13_=3.1, p=0.1028; Figure 1G-I; Figure 1 – figure supplement C-D). However, an overall decrease in exploratory activity in the light zone was observed (OFF-ON-OFF, F_2,30_=37.8, *p*<0.0001). In the NSF, no significant differences in the latency to reach the center or latency to feed were observed (Figure 1J-M).

Importantly, D1-VP-ChR2 mice did not show differences in time spent or distance travelled in the OF arena in comparison with D1-VP-YFP mice (Figure 1 – figure supplement E-H), suggesting that the optical stimulation protocol adopted in this study caused no gross locomotor alterations.

These data show that acute optical activation of D1-MSN-VP projections causes a *very mild* anxious-like behavior.

### Optical activation of D1-MSN projections to the VTA does not alter anxious-like behavior

To explore whether D1-MSN-VTA projections play a role in anxious-like behaviors, we used a similar strategy as above, but the optical fiber was placed in the VTA (Figure 2A). About 25% of the VTA area presented YFP innervation (D1-MSN terminals) (Figure 2B-C).

**Figure 2.**
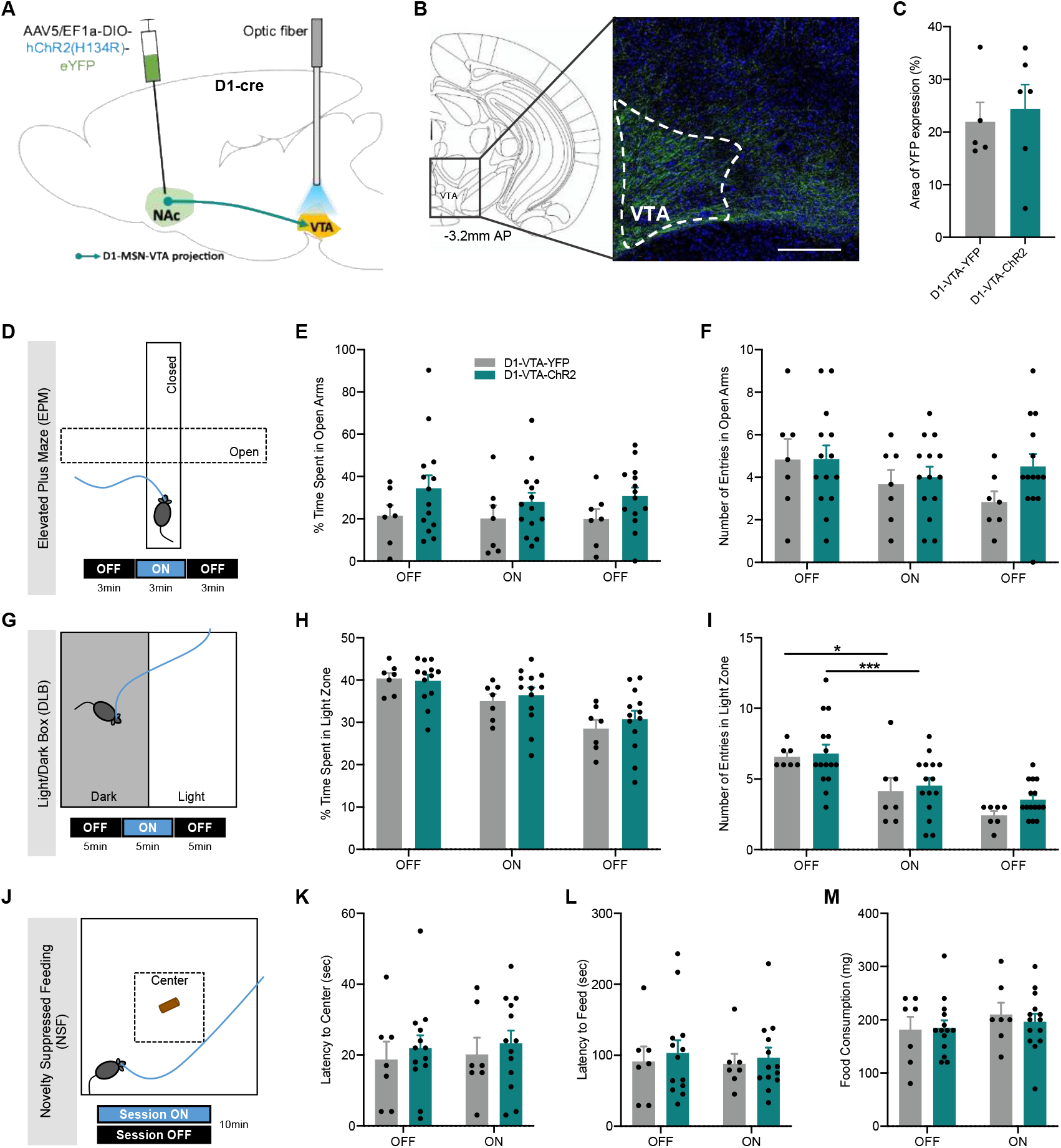
Optogenetic stimulation of D1-MSNs projecting to the VTA does not induce anxious-like behavior. **A** NAc D1-MSNs of D1-cre mice were injected with either cre-dependent ChR2 or YFP. Blue light was delivered through an optical fiber implanted in the VTA to allow terminal stimulation (473nm, 20Hz, 25ms pulses, 50% duty cycle). **B** Coronal slice showing expression of YFP using immunostaining. Numbers represent distance to bregma in millimeters; AP: anteroposterior; scale bar: 500 μm. **C** About 25% of VP region presented YFP expression, indicative of D1-MSN terminals. **D** The EPM was divided in 3-min epochs: OFF-ON-OFF. Optical stimulation of D1-MSN-VTA did not change **E** the time spent or **F** the number of entries in the open arms. **G** The LDB test was divided in 5-min epochs: OFF-ON-OFF. No differences were observed in the **H** time spent or **I** number of entries in the light zone of the arena. **J** Two sessions of NSF were performed: 1 session ON and 1 session OFF (counterbalanced); optical stimulation was performed throughout the entire session ON. D1-VTA-ChR2 mice showed a similar **K** latency to reach the food pellet located in the center of the arena and **L** latency to feed in the ON and OFF sessions. **M** food consumption after the NSF was not altered. Data denote mean±SEM. *p≤0.05, **p≤0.01, ***p≤0.001.

Importantly, no changes in anxiety-like behaviors were observed with optical stimulation of D1-MSN-VTA in the EPM (Figure 2D-F; Figure 2 – figure supplement A-B). In the LDB, although no differences were observed between experimental groups, a significant decrease in overall exploration of the apparatus was observed (time spent in light zone: F_1,12_=0.2, p=0.6914; number of entries in light zone: F_1,14_=1.4, p=0.2589; Figure 2G-I; Figure 2 – figure supplement C-D). In the NSF, no effect of stimulation was observed (Figure 2J-M).

D1-VTA-ChR2 mice showed no changes in locomotion as measured by the OF (Figure 2 – figure supplement E-H).

These data show that acute optical activation of D1-MSN-VTA projections did not impact anxious-like behaviors.

### Optical activation of D2-MSN-VP projections drives anxious-like behavior

Next, to determine the influence of D2-MSNs projecting to the VP in the induction of anxious-like behavior, we unilaterally injected D2-cre mice in the NAc with a cre-dependent ChR2 in fusion with YFP, or the control YFP virus (Figure 3A); an optical fiber was placed in the VP to allow optical activation of D2-MSN terminals. Nearly 25% of the VP area presented YFP innervation (D2-MSN terminals) (Figure 3B-C).

**Figure 3.**
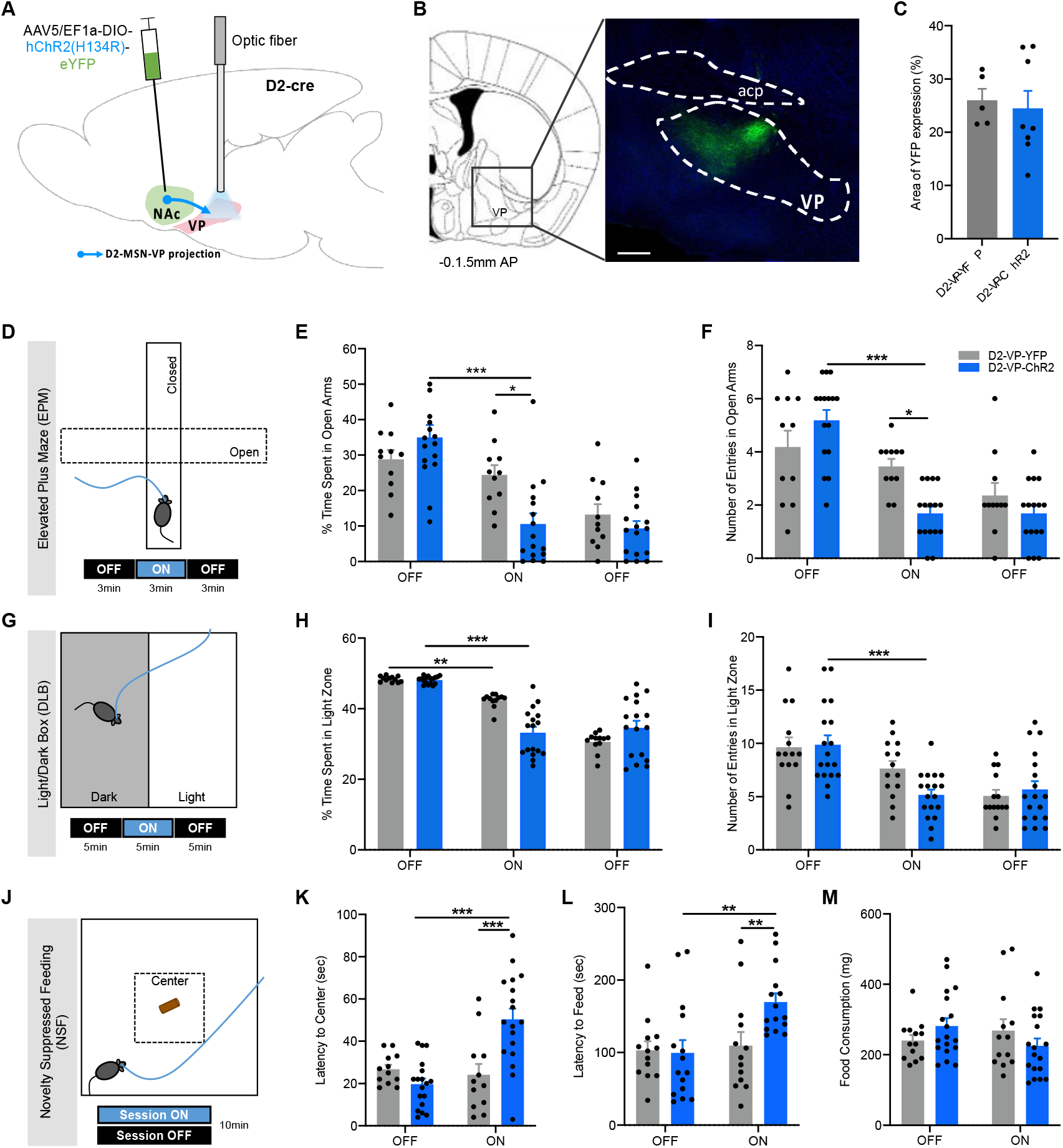
Optogenetic stimulation of D2-MSNs projecting to the VP triggers anxious-like behavior. **A** NAc D2-MSNs of D2-cre mice were injected with either cre-dependent ChR2 or YFP. Blue light was delivered through an optical fiber implanted in the VP (473nm, 20Hz, 25ms pulses, 50% duty cycle). **B** Coronal slice showing expression of YFP using immunostaining. Numbers represent distance to bregma in millimeters; AP: anteroposterior; scale bar: 500 μm. **C** About 25% of VP presented YFP expression, indicative of D2-MSN terminals. **D** The EPM was divided in 3-min epochs: OFF-ON-OFF. D2-MSN-VP optical stimulation significantly decreased **E** the time spent and **F** the number of entries in the open arms of the EPM. **G** The LDB test was divided in 5-min epochs: OFF-ON-OFF. **H** D2-VP-ChR2 significantly decreased the time spent in the light zone of the arena in the ON period but not the **I** number of entries. **J** Two sessions of NSF were performed: 1 session ON and 1 session OFF (counterbalanced); optical stimulation was performed throughout the session ON. D2-VP-ChR2 mice showed a significant higher **K** latency to reach and to **L** consume the pellet in comparison to YFP mice and to the OFF session. **M** No changes in food consumption after the NSF. Data denote mean±SEM. *p≤0.05, **p≤0.01, ***p≤0.001.

Interestingly, optical stimulation of D2-MSN-VP projections significantly decreased the time spent in the open arms of the EPM (F_2,36_=39.1, *p*<0.0001; OFF *vs* ON, *p*<0.0001; ChR2 *vs* YFP, *p*=0.0412; Figure 3D, E; Figure 3 – figure supplement 3A), and decreased the number of entries in the open arms (F_2,34_=30.5, *p*<0.0001; OFF vs ON, p<0.0001; ChR2 *vs* eYFP, *p*=0.050; Figure 3F; Figure 3 – figure supplement 3B). These data are supported by the LDB results, since ChR2 mice also showed decreased time spent in the light zone of the arena (F_2,32_=120.3, *p*<0.0001; OFF vs ON, *p*<0.0001; ChR2 vs YFP in ON epoch, *p*=0.0002; Figure 3G-H; Figure 3 – figure supplement C;). The number of entries in the light zone was similar between groups (Figure 3I-J; Figure 3 – figure supplement 3D).

In further support of an anxiogenic role of D2-MSN-VP projections, D2-MSN-VP-stimulated mice showed a significant increase in the latency to reach the center (F_1,28_=13.9, p=0.0009; OFF vs ON, *p*<0.0001; Figure 3J-K) and to initiate feeding (F_1,26_=8.0, p=0.0089; OFF vs ON, *p*=0.0016; Figure 3l) in the NSF session receiving optical stimulation in comparison with the OFF session. In the ON session, D2-VP-ChR2 mice presented a significant increase in comparison with D2-VP-YFP control group in both latency to reach the center and latency to initiate a feeding episode in the center of the arena (D2-VP-ChR2 vs D2-VP-YFP; latency to center: F_1,28_=4.5, *p*=0.0439, ChR2 vs YFP in ON epoch, *p*=0.0012; latency to feed: F_1,26_=2.7, p=0.1123, ChR2 vs YFP in ON epoch, *p*=0.0167). No differences in food consumption after the NSF test were observed (Figure 3M).

Importantly, optical activation of D2-MSN-VP projections does not induce locomotor alterations in the OF (Figure 3 – figure supplement E-H).

These data show that acute optical activation of D2-MSN-VP projections triggers an anxious-like behavior in different behavioral tests.

### Electrophysiological correlates of D1-MSN stimulation

Next, we performed extracellular single-cell electrophysiological recordings in anaesthetized mice previously injected with a cre-dependent ChR2 virus in the NAc of D1-cre mice (Figure 4A-B). A recording electrode glued to an optical fiber was descended to the target region - NAc for cell somas, VP or VTA for terminal-evoked activity. We applied the same OFF-ON-OFF epoch stimulation protocol that was used during behavior (3min OFF; 3 min ON (20Hz, 25ms light pulses, 50% duty cycle, 5mW at the tip off the fiber); 3 min OFF). In the NAc, we observed increased neuronal activity in the range of *physiological* levels with this stimulation parameter (Figure 4 – figure supplement 1 A; change in firing rate from ~2Hz to ~4.6Hz at the ON epoch).

**Figure 4.**
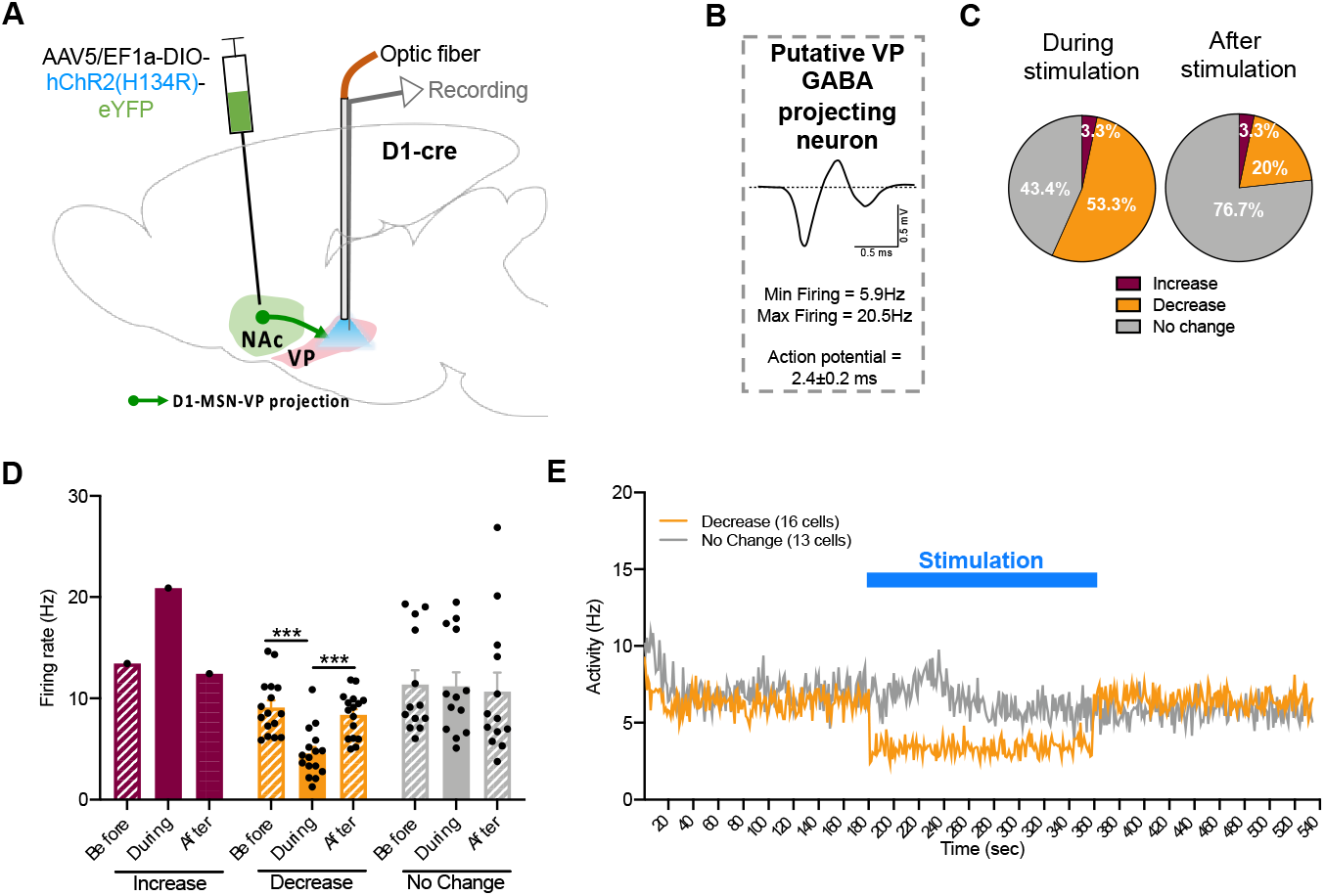
VP electrophysiological responses to optical stimulation of D1-MSN-VP projections. **A** Schematic representation of VP electrophysiological recordings. **B** Representative waveform of a VP putative GABAergic neuron. **C** Pie charts represent the percentage of each type of response to stimulus, showing a high percentage of VP putative GABAergic neurons decreasing activity during stimulation. **D** Bar graphs represent net firing rate before, during and after optogenetic stimulation. **E** Temporal variation of VP activity; note the decrease in activity during optical stimulation (blue). Data denote mean±SEM. *** p<0.001.

Next, we evaluated the neuronal activity of putative GABAergic projecting neurons of the VP (Pang et al., 1998) in response to D1-MSN terminal stimulation (Figure 4A-B). Optical stimulation of D1-MSN-VP projections significantly decreased the average firing rate of 53.3% of identified VP putative GABAergic projecting neurons (decrease: F_2,15_=35.2, *p*<0.0001; OFF vs ON *p*<0.0001; Figure 4C-E); 3.3% increased activity; 43.4% of recorded neurons did not change activity.

Next, we also evaluated the neuronal activity of putative dopaminergic and GABAergic neurons of the VTA (Totah et al., 2013; Ungless, 2004) (Figure 5A-B). Optical stimulation of D1-MSN-VTA projections significantly decreased the average firing rate of 59% of VTA putative dopaminergic neurons (24/39 neurons; F_2,22_=38.6, *p*<0.0001; OFF vs ON *p*<0.0001) and of 60% of putative GABAergic neurons (3/5 neurons; Figure 5C-E), indicating an overall decreased activity of the VTA.

**Figure 5.**
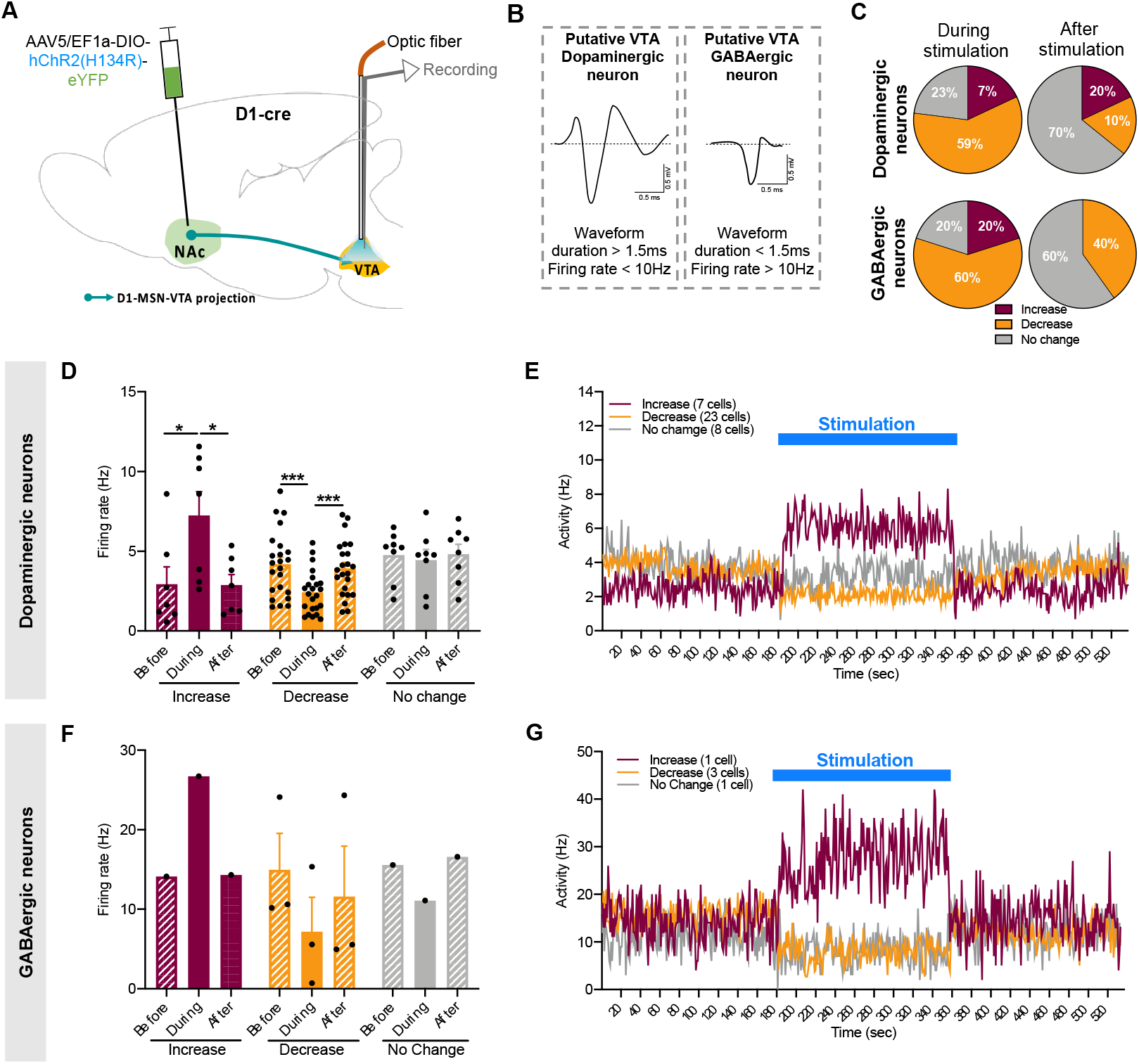
VTA electrophysiological responses to optical stimulation of D1-MSN-VTA projections. **A** Schematic representation of VTA electrophysiological recordings. **B** Representative waveform of a VTA putative dopaminergic neuron and a putative GABAergic neuron. **C** Pie charts represent the percentage of each type of response to stimulus, showing a significant percentage of VTA putative dopaminergic and GABAergic neurons changing activity during stimulation. Bar graphs represent net firing rate before, during and after optogenetic stimulation of putative **D** dopaminergic neurons and **F** putative GABAergic neurons. Temporal variation of activity of VTA **E** putative dopaminergic neurons and **G** putative GABAergic neurons. Data denote mean±SEM. * p<0.05; ** p<0.001.

These data demonstrate that D1-MSN optical manipulation significantly alters NAc, VP and VTA neuronal activity, though it did not trigger major changes in behavior.

### Electrophysiological correlates of D2-MSN stimulation

Our behavioral results show that optogenetic stimulation of D2-MSN-VP projections triggered an anxious-like phenotype. We performed extracellular single-cell electrophysiological recordings in anaesthetized mice previously injected with a cre-dependent ChR2 virus in the NAc of D2-cre mice (Figure 6A-B).

**Figure 6.**
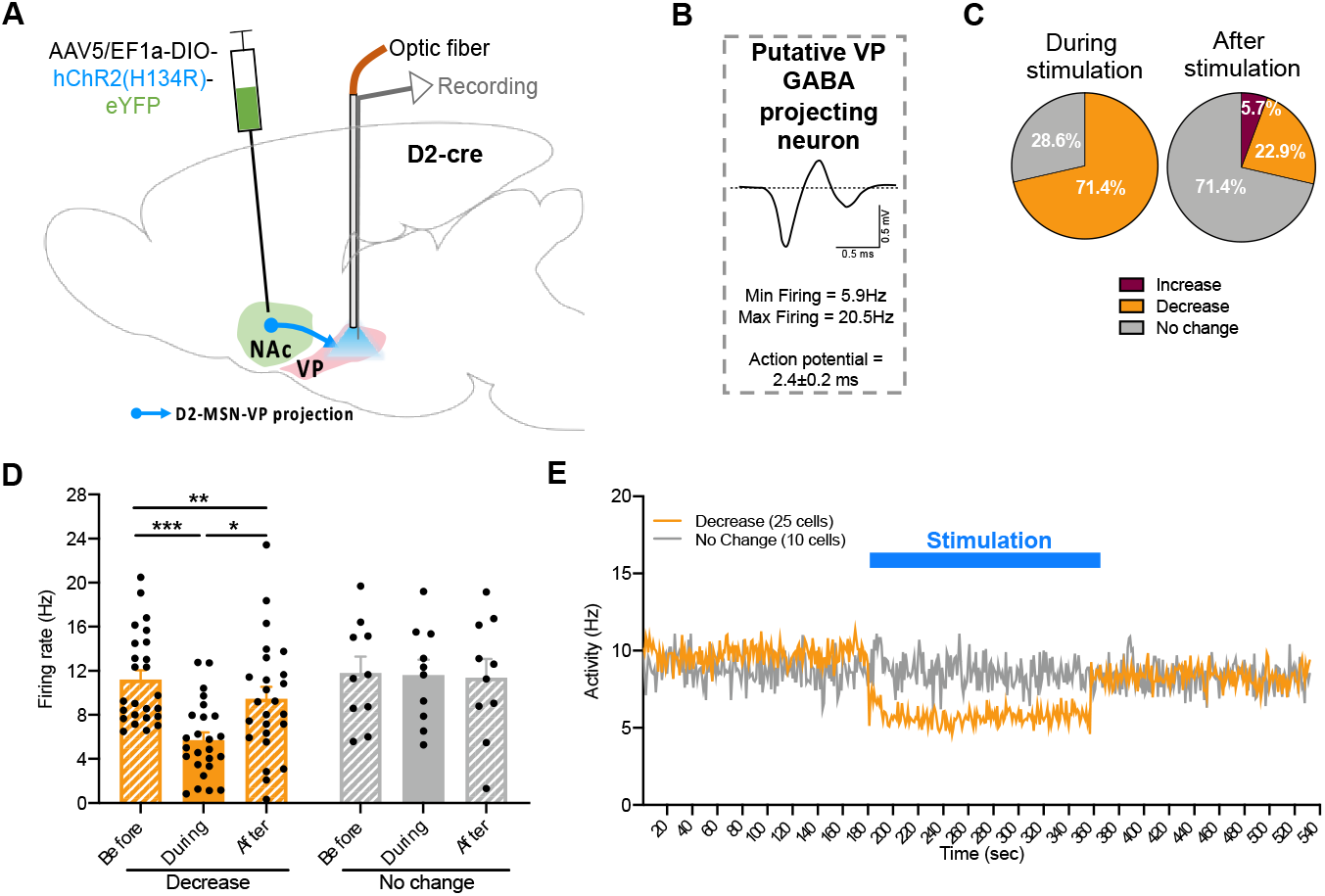
VP electrophysiological responses to optical stimulation of D2-MSN-VP projections. **A** Schematic representation of VP electrophysiological recordings. **B** Representative waveform of a VP putative GABAergic neuron. **C** Pie charts represent the percentage of each type of response to stimulus, showing a high percentage of VP putative GABAergic neurons decreasing activity during stimulation. **D** Bar graphs represent net firing rate before, during and after optogenetic stimulation. E Temporal variation of VP activity; note the decrease in activity during optical stimulation (blue). Data denote mean±SEM. * p<0.05; ** p<0.01; *** p<0.001.

In the NAc, we observed increased neuronal activity in the range of *physiological* levels with this stimulation parameter (Figure 4 – figure supplement 1 B; change in firing rate from ~2Hz to ~4.6Hz at the ON epoch).

Optical stimulation of D2-MSN-VP projections significantly decreased the average firing rate of 71.4% of VP putative GABAergic projecting neurons (25/35 neurons; F_2,24_=16.7, *p*<0.0001; OFF vs ON *p*=0.0001; Figure 6C-E), while 28.6% of recorded neurons did not change their activity.

These data indicate that D2-MSN manipulation triggered more evident changes in VP GABAergic activity than D1-MSN manipulation, which may support the anxiogenic effect observed for D2-MSNs but not D1-MSNs manipulation.

### Pre-treatment with diazepam prevents D2-MSN-VP-induced anxious-like phenotype

The preceding results showed an anxious-like behavior triggered by D2-MSN-VP stimulation and a GABA-mediated effect of D2-MSN terminal stimulation in the VP, leading to reduced VP net activity. So, we decided to treat animals systemically with a low dose of a conventional anxiolytic drug, diazepam, a positive allosteric modulator of the GABA type A receptors (Löw et al., 2000). We used a dosage that was previously shown to not have a significant anxiolytic effect under physiological conditions (Pádua-Reis et al., 2021). For this experiment, we injected a new set of D2-cre mice with ChR2 (or YFP in control group) in the NAc and implanted an optical fiber in the VP to allow optical stimulation. 30 minutes before behavioral testing, mice received an intraperitoneal (i.p.) injection of diazepam at 0.5mg/kg (DIA), or vehicle (VEH) (Fig. 7A).

**Figure 7.**
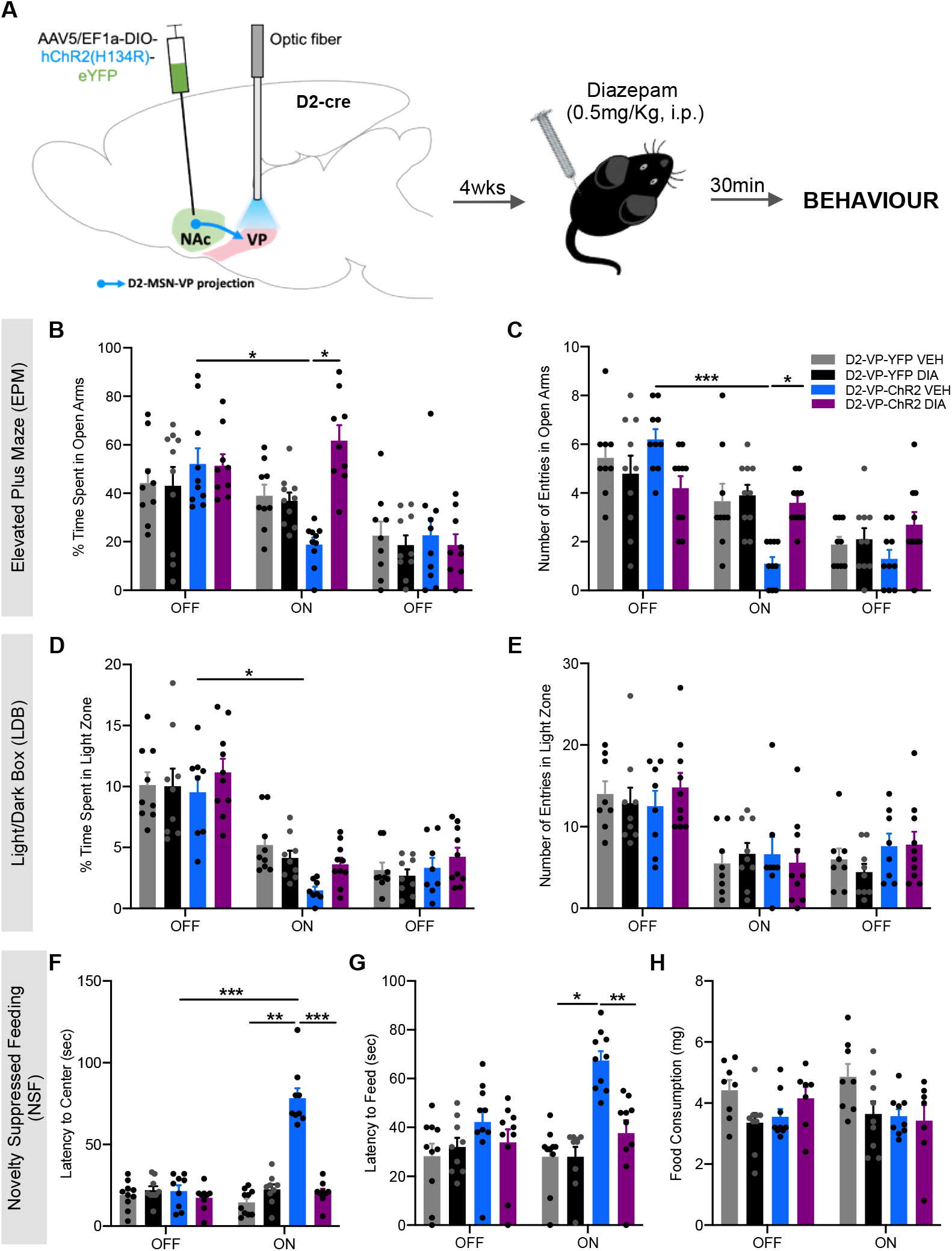
Pre-treatment with diazepam prevents the anxious-like phenotype induced by optogenetic stimulation of D2-MSN-VP terminals. **A** D2-cre mice were transfected with a cre-dependent ChR2 or YFP in controls, in the NAc and an optical fibre was implanted in the VP to allow D2-MSN terminal stimulation. 4 weeks after, mice were injected with diazepam (DIA; 0.5mg/kg i.p.) or vehicle (VEH) 30 min before behavioral evaluation - EPM, LDB test, NSF test and OF test. Behavioral tasks were performed 72h apart from each other. Diazepam (DIA) treatment significantly decreased **B** the time spent and **C** the number of entries in the open arms of the EPM of ChR2 mice. Similarly, in the LDB test, diazepam treatment reverted the decrease in **D** the time spent in the light zone of the arena, leaving **E** the number of entries unaltered. D2-ChR2-DIA mice also reverted the anxious phenotype of the NSF test, presenting a significant decrease in the latency to **F** reach the center and to **G** consume the food pellet. **H** no changes in food consumption after the NSF. In all behavioral tests performed, diazepam treatment did not change basal anxiety states in YFP mice. Data denote mean±SEM. * p<0.05; ** p<0.01; *** p<0.001.

In the EPM, no effect of diazepam was found in control YFP mice (Figure 7). D2-VP-ChR2 VEH mice showed a significant decrease in the time spent in open arms (F_2,18_=29.5, *p*<0.0001; baseline vs ON, *p*=0.0002; Figure 7B; Figure 7 – figure supplement 1 A-B), and decreased number of entries in the open arms (Figure 7C; F_2,18_=55.8, *p*<0.0001; baseline vs ON, *p*<0.0001) during the optical stimulation period (ON), confirming here-reported anxiogenic effect. Conversely, D2-VP-ChR2 DIA mice spent a significantly higher amount of time in open arms (ChR2 DIA vs ChR2 VEH, *p*<0.0001) during stimulation in comparison with D2-VP-ChR2 VEH animals.

In the LDB test, D2-MSN-VP optical activation induced an anxious-like behavior (F_2,18_=88.1, p<0.0001, ChR2 VEH OFF vs ChR2 VEH ON, p<0.0001) that was no longer observed after treatment with diazepam (Figure 7D-E; Figure 7 – figure supplement 1C-D).

In the NSF, treatment with diazepam prevented the anxious phenotype caused by optical stimulation of D2-MSN-VP projections. Importantly, in the ON session, D2-VP-ChR2 DIA mice displayed a significant decrease in the latency to reach the center of the arena (Figure 7H; F_1,9_=49.1, *p*<0.0001; ChR2 DIA vs ChR2 VEH, *p*<0.0001) and latency to initiate feeding (Figure 7I; F_1,9_=7.1, *p*=0.0253; ChR2 DIA vs ChR2 VEH, *p*<0.0001) in comparison with D2-VP-ChR2 VEH, and comparable to D2-VP-YPF VEH control mice. No effect of diazepam was found in control YFP animals. Food consumption after the NSF test was similar between all groups (Figure 7J).

Importantly, no sedative effect of diazepam was found in the OF test (Figure 7 – figure supplement 1F-I).

Overall, these results indicate that diazepam treatment prevented the development of the anxiogenic phenotype caused by acute D2-MSN-VP optical stimulation, with no impact in control animals.

## Discussion

In this work we showed that acute optogenetic activation of NAc D2-MSN-VP projections elicits a transient anxious-like phenotype. D2-MSN optical activation triggered a decrease in VP GABAergic activity. We further show that modulating GABAergic activity through the systemic administration of the anxiolytic drug diazepam, prevented the anxious state triggered by stimulation of D2-MSN-VP projections. Manipulation of NAc D1-MSNs to VP results in very mild anxious-like behavioral effect in some tests, while D1-MSN to VTA had no observable effects in anxious-like behavior.

There is sparse human evidence linking NAc to anxiety. Neuroimaging studies have shown that patients suffering from social anxiety disorder have increased activity in the NAc (Kilts et al., 2006). Also, in an avoidance task, accumbal activation was associated with anxiety behavior (Levita et al., 2012). Interestingly, deep brain stimulation (DBS) of NAc has shown to be effective in reducing anxiety in depressed patients (Bewernick et al., 2012). Regarding animal models, it is obvious that present inherent limitations for the study of psychiatric conditions, but since mensurable behavioral readouts indicative of anxious-like phenotype exist, these models are valuable in the study of the mechanisms underlying anxiety. Anxiety disorders are classically maintained by maladaptive approach-avoidance conflict behaviors (American Psychiatric Association, 2013). Avoidance can be considered a motivated behavior in response to stimuli that threaten the individual’s well-being (e.g., fear or pain), while approach can be considered a drive motivated by stimuli/events that ensure well-being (e.g., rewarding). Importantly, these features can be assessed in rodents through behavioral tests, namely the EPM, LDB and NSF tests (Felix-Ortiz et al., 2016; Kim et al., 2013), as they rely on the premise that animals with an anxious-like phenotype will avoid threatening/unsafe open and bright spaces. To date, the available studies evaluated the contribution of NAc MSNs in animal models already presenting anxious-like phenotype. However, in this work, we modulated NAc activity in naïve animals in order to observe if an anxious-like phenotype could be elicited.

Acute D1-MSN-VP activation triggered a mild anxious-like phenotype in the EPM test, but not in the other anxiety tests. Interestingly, we observe a similar trend for decreased activity of both dopaminergic and GABAergic VTA neurons due to D1-MSN-VTA activation. A recent study showed that chronic activation of VTA dopaminergic neurons and inhibition of NAc medial shell neurons projecting to VTA dopaminergic neurons triggers anxiety (Qi et al., 2022).

Acute D2-MSN-VP input activation triggered a more evident anxious-like phenotype in the EPM, LDB and NSF tests. D2-MSN-VP activation was associated with a concomitant decrease in VP GABAergic activity, consistent with the inhibitory (GABAergic) nature of D2-MSNs. Our data is in agreement with other studies showing that inhibition of VP GABA neurons induces place aversion and a sense of threat, and increases defense behavior (Moaddab et al., 2021; Wulff et al., 2019). Importantly, the decrease in VP GABAergic activity here observed may lead to increased VTA GABAergic activity (Soares-Cunha et al., 2020, 2018), suppressing VTA dopaminergic neurons to promote a generalized anxiety-like phenotype (Zweifel et al., 2011).

Given that the data pointed to changes in VP GABA neurons, we administered a low dose of diazepam 30 minutes before behavioral assessment with optogenetic activation of D2-MSN-VP inputs. This pre-treatment prevented the anxious-like behavior caused by optogenetic activation of D2-MSN-VP inputs. Benzodiazepines induce hyperpolarization of VTA GABAergic interneurons (Tan et al., 2012), likely disinhibiting VTA dopamine neurons (O’Brien et al., 1998). This evidence further supports our hypothesis that optogenetic activation of D2-MSN-VP projections can indirectly lead to blunted dopaminergic activity, and consequently induce an anxious state (Zweifel et al., 2011). It is important to refer that we selected a low dose of diazepam that did not exert an observable effect in control animals (Pádua-Reis et al., 2021), because we aimed to abolish optically-induced GABA-mediated changes in stimulated animals, rather than exerting a robust and *unspecific/general* anxiolytic effect.

In this work we adopted a strategy that would allow assess anxiety-like behavioral effects of stimulation within the same animal, by including OFF and ON epochs, whenever possible, in the behavioral tasks. Several others have used the same approach to show that the amygdala, the prefrontal cortex and the VTA can modulate anxiety-like behaviors (Felix-Ortiz et al., 2016; Kim et al., 2013; Qi et al., 2022). However, it is important to refer that we observed an overall decrease in exploratory behavior throughout the tests, which hampers some of the analyses.

The optical stimulation parameters chosen, which are also similar to the ones of other studies (Felix-Ortiz et al., 2016; Kim et al., 2013), significantly increased accumbal neuronal activity within the range of what is observed *in vivo* under physiological conditions in animals engaged in learning tasks (Cerri et al., 2014; West and Carelli, 2016). In addition, both D1- and D2-MSN terminal activation resulted in significant changes in downstream targets, VP and VTA, which was also previously shown to trigger rewarding/aversive conditioning (Soares-Cunha et al., 2022, 2020).

Though there are limitations in directly confirming electrophysiological recordings in different animals, it was interesting to observe that D2-MSN-VP manipulation triggered more evident neuronal activity changes in the VP than D1-MSN-VP activation (with 75% *versus* 53% decreased VP GABAergic activity). Nevertheless, This data is in agreement with the fact that VP neurons receive higher innervation from D2-MSNs than from D1-MSNs (Kupchik et al., 2015).

In sum, we show that optical activation of NAc D2-MSN-VP inputs triggers a transient anxious-like phenotype that can be prevented by diazepam pre-treatment. These findings highlight the need to perform additional studies to investigate how distinct subpopulations of NAc neurons contribute for the development of anxious-like behaviors.

## Material and Methods

### Subjects

C57BL6/J transgenic male heterozygous D1-cre (Gensat, EY262) and D2-cre (Gensat, ER44) mice (2-4 months old at the beginning of the experiments) were used. Animals were housed in groups of 3-5 animals and were maintained under standard laboratory conditions: 12h light/dark cycle (lights on at 8:00am), temperature of 22°C±1°C and 60% relative humidity; standard diet and water *ad libitum* were provided, except when stated otherwise. All behavioral experiments were performed during the light period of the light/dark cycle (between 8:30 am and 2:30pm).

Health monitoring was performed according to FELASA guidelines. All animal experiments complied to the ARRIVE guidelines and were conducted in accordance to European Union Regulations (Directive 2010/63/EU). Animal facilities and experimenters were certified by the Portuguese regulatory entity, Direção-Geral de Alimentação e Veterinária (DGAV). All protocols were approved by the Ethics Committee of the Life and Health Sciences Research Institute and by DGAV (protocol #19074).

### Genotyping

DNA was isolated from tail biopsy using Citogene DNA isolation kit (Citomed, Lisbon, Portugal). In a single PCR genotyping tube, the primers Drd1a F1 (5’-GCTATGGAGATGCTCCTGATGGAA-3’) and CreGS R1 (5’-CGGCAAACGGACAGAAGCATT-3’) were used to amplify the D1-cre transgene (340 bp), and the primers Drd2 F1 (5’-GTGCGTCAGCATTTGGAGCAA-3’) and CreGS R1 (5’- CGGCAAACGGACAGAAGCATT-3’) were used to amplify the D2-cre transgene (700 bp). An internal control gene (lipocalin, 500 bp) was used in the PCR (LCN_1 (5’- GTC CTT CTC ACT TTG ACA GAA GTC AGG −3’) and LCN_2 (5’- CAC ATC TCA TGC TGC TCA GAT AGC CAC −3’)). Heterozygous mice were differentiated from the wild-type mice by the presence of two amplified DNA products corresponding to the internal control gene and the transgene. Gels were visualized with GEL DOC EZ imager (Bio-Rad, Hercules, CA, USA) and analyzed with Image Lab 4.1 (Bio-Rad, Hercules, CA, USA).

### Virus constructs

Cre-inducible AAV5/EF1a-DIO-hChR2(H134R)-eYFP and AAV5/EF1a-DIO-eYFP viruses were obtained directly from the UNC Gene Therapy Center (University of North Carolina, NC, USA). AAV5 vectors titers were 3.7–6 × 10^12^ viral molecules/ml as determined by dot blot.

### Virus injection and fiber implantation

D1-cre and D2-cre heterozygous transgenic male mice were submitted to stereotaxic surgeries. Mice were anaesthetized with 75mg/kg ketamine (Imalgene, Merial, Lyon, France) plus 1mg/kg medetomidine (Dorbene, Cymedica, Horovice, Czech Republic). 500nl of a cre-inducible AAV5/EF1a-DIO-hChR2(H134R)-eYFP virus (or control AAV5/EF1a-DIO-eYFP) was unilaterally injected in the right hemisphere of the NAc (coordinates (Paxinos and Franklin, n.d.): +1.3mm anteroposterior (AP), +0.9mm mediolateral (ML), and −4.0mm dorsoventral (DV)) using a 30-gauge needle Hamilton syringe (Hamilton Company, Reno, NV, USA), at a rate of 100nl/min. After injection, the syringe was left in place for 5 minutes to allow viral diffusion. An optical fiber (200μm core fiberoptic; Thorlabs, Newton, NJ, USA) with 2.5mm stainless steel ferrule (Thorlabs, Newton, NJ, USA) was unilaterally implanted in the right hemisphere of the VP (for D1- and D2-cre mice; coordinates (Paxinos and Franklin, n.d.): −0.1mm AP, +1.6mm ML, −3.9mm DV) or in the right hemisphere of the VTA (for D1-cre mice; coordinates (Paxinos and Franklin, n.d.): −3.2mm AP, +0.5mm ML, −4.5mm DV), and was secured to the skull with dental cement (C&B kit, Sun Medical, Shiga, Japan), to allow optogenetic stimulation of D1- or D2-MSN terminals in downstream targets. After the surgery, mice were given an anesthetic reversal – 1mg/kg atipamezole (Antisedan, Zoetis, Parsippany, NJ, USA), and let to recover for four weeks before initiation of the behavioral procedures.

Animals assigned for the electrophysiological experiment were not implanted with the optical fiber. After viral injection, animals were removed from the stereotaxic frame, were sutures, were given an anesthetic reversal – 1mg/kg atipamezole (Antisedan, Zoetis, Parsippany, NJ, USA), and let to recover for four weeks before the electrophysiological recordings.

All animals were treated 30 minutes before surgery and 6hours after surgery with an analgesic – 0.1mg/kg buprenorphine (Bupaq, Richter Pharma, Wels, Austria). The same treatment was maintained once a day for three consecutive days after surgery. If animals displayed severe loss of body weight a multivitamin supplement would also be administered.

### Optogenetic manipulation

Optogenetic stimulation was performed using 5mW blue light generated by a 473nm DPSS laser (CNI, Changchun, China) and a pulse generator (AMPI, MN, USA), delivered through an optic fiber cable (0.22NA, 200μm diameter; Thorlabs, NJ, USA) attached to the implanted ferrule. Laser output and pulsed light were controlled by a pulse generator (Master-8; AMPI, New Ulm, MN, USA). The stimulation was performed as follows: 25ms light pulses at 20Hz, 50% duty cycle; the duration of stimulation varied with behavioral protocol (3- or 5-min epochs).

### Drug

Diazepam (DIA; 0.5mg/kg; Labesfal, Tondela, Portugal) and vehicle (VEH; 0.9%saline) were injected intraperitoneally (i.p.) 30min before the behavioral tests. Drug doses were based on a previous study (Pádua-Reis et al., 2021)

### Experimental cohorts

For behavioral experiments with optogenetic stimulation, transgenic male mice were randomly distributed into different groups depending on the transgenic line: D1-VP-ChR2 (n = 16); D1-VP-YFP (n = 7); D1-VTA-ChR2 (n = 15); D1-VTA-YFP (n = 7); D2-VP-ChR2 (n = 19); D2-VP-YFP (n = 13).

For drug experiment, transgenic D2-cre mice were randomly divided into different groups: D2-VP-YFP VEH (n = 9); D2-VP-YFP DIA (n = 10); D2-VP-ChR2 VEH (n = 10); D2-VP-ChR2 DIA (n = 10).

For the *in vivo* electrophysiological experiment, 15 transgenic D1-cre and 14 transgenic D2-cre mice with 6 months of age were used.

### Behavioral Assessment

Behavioral tests started four weeks after surgery. All behavioral tests were performed and analyzed blindly. Mice were transferred to the testing rooms 30min before the beginning of each test to allow acclimation. All animals performed all the behavioral tests, in the following order: behavioral test day 1 – elevated plus maze; behavioral test day 4 – light/dark box test; behavioral test day 6 – open field test; behavioral test day 8 – novelty suppressed feeding (session 1); behavioral test day 14 – novelty suppressed feeding (session 2).

### Elevated Plus Maze (EPM)

The EPM test was adapted from a previously described protocol (Felix-Ortiz et al., 2016). The EPM apparatus consisted of a black acrylic plus-shaped platform with two open and illuminated arms (50.8×10.22×40.6cm) and two enclosed and dark arms (50.8×10.2×40.6cm) elevated 72.4cm of the floor (Med Associates Inc., St. Albans, VT, USA). Individual mice were attached to the optical fiber cable and placed longitudinally in the central platform of the EPM apparatus (Med Associates Inc., VT, USA), always facing the same corner. Each session had a duration of 9min, divided in three alternating 3-min epochs: no laser stimulation, laser stimulation, no laser stimulation (OFF-ON-OFF). All sessions were recorded by video and analyzed in the EthoVisionX13 software (Noldus, the Netherlands). Time spent in open and closed arms, and number of entries in the open and closed arms were assessed. Mice showing absence of exploratory behavior to the open arms in the first OFF epoch (0% time spent and 0 entries in open arms) were excluded from the analysis of the behavioral test: 4 D1-VP-ChR2, 1 D1-VTA-ChR2, 1 D2-VP-ChR2 and 2 D2-VP-YFP mice were removed.

### Open Field (OF)

The OF test was adapted from a previously described protocol (Felix-Ortiz et al., 2016). The OF arena consisted of a square white floor (43.2cmx43.2cm) with four transparent acrylic walls (Med Associates Inc., St. Albans, VT, USA). Individual mice were attached to the optical fiber cable and placed in the center of the arena (Med Associates Inc., VT, USA), always facing the same wall. Each session had a duration of 9min, divided in three alternating 3-min epochs: OFF-ON-OFF.

Distance travelled, number of entries and time spent in each zone were measured using an automated video-tracking system (Activity Monitor software; Med Associates Inc., St Albans, VT, USA). Mice exhibiting absence of exploratory behavior to the center of the open field arena in the first OFF epoch (0% time spent and 0 entries in center of the arena) were excluded from the analysis of the behavioral test: 2 D1-VP-ChR2, 3 D1-VTA-ChR2 and 1 D2-VP-YFP mice were removed.

### Light/Dark Box (LDB) test

The LDB was adapted from a previously described protocol (Kim et al., 2013). The LDB apparatus consisted of an arena similar to the one used in the OF test (43.2cmx43.2cm; Med Associates Inc., St. Albans, VT, USA), divided half-dark and half-light using an acrylic chamber. Individual mice were attached to the optical fiber cable and placed in the center of the dark compartment. Each session had a duration of 15min, divided in three alternating 5-min epochs: OFF-ON-OFF.

Distance travelled, number of entries and time spent in each zone were measured using an automated video-tracking system (Activity Monitor software; Med Associates Inc., St Albans, VT, USA). Mice exhibiting absence of exploratory behavior to the light compartment of the box in the first OFF epoch (0% time spent and 0 entries in light side) were excluded from the analysis of the behavioral test: 2 D1-VTA-ChR2 and 2 D2-VP-YFP mice were removed.

### Novelty Suppressed Feeding (NSF)

The NSF was adapted from a previously described protocol (Felix-Ortiz et al., 2016). The apparatus used for the test was similar to the OF arena (Med Associates Inc., VT, USA) with the floor covered with clean corn cob bedding. One familiar food pellet (Mucedola 4RF21-GLP) weighing approximately 6g was placed in the center of the arena. Mice food-deprived for 18h were attached to the optical fiber cable and placed in one corner of the arena. The latency to reach the food pellet and the latency to begin a feeding episode were manually recorded. The session ended when mice first fed or after 10min elapsed without consumption. Immediately after testing, mice were removed from the arena and were individually housed, and free food consumption was measured after 30min. The task was repeated one week after, counterbalanced for laser stimulation session (ON) or no stimulation session (OFF). In the ON session, stimulation was performed for all the period until the animal first fed. Mice showing absence of exploratory behavior during the 10min session (no exploration to the center of the arena or the food pellet) were removed from the analysis of the behavioral test. 1 D1-VP-ChR2, 1 D1-VP-YFP, 1 D1-VTA-ChR2, 1 D1-VTA-YFP and 5 D2-VP-ChR2 mice were removed.

### *In vivo* single-cell electrophysiology recordings

Mice were anaesthetized with urethane (1.75g/kg; Sigma now Merck KGaA, Darmstadt, Germany), divided into 3 doses administered intraperitoneally with an interval of 30 minutes. Mice were placed in the stereotaxic frame (David Kopf Instruments, Tujunga, CA, USA) with nontraumatic ear bars (Stoelting, Wood Dale, IL, USA). A tungsten recording electrode (tip impedance 5-10Mat 1kHz) coupled with an optical fiber cable (Thorlabs, Newton, USA) was placed in the VP (D1-VP-ChR2 and D2-VP-ChR2 mice; coordinates (Paxinos and Franklin, n.d.): −0.1mm AP, +1.6mm ML, - 3.5 to −4.5mm DV), or in the VTA (D1-VTA-ChR2 mice; coordinates (Paxinos and Franklin, n.d.): - 3.2mm AP, +0.5mm ML, −4.0 to −4.8mm DV). A ground screw was placed in the skull to close the circuit allowing signal stabilization. The recordings were performed under a Faraday box to reduce background.

Single neuronal activity was recorded extracellularly, and recordings were amplified and filtered by the Neurolog amplifier (NL900D, Digitimer Ltd, Hertfordshire, UK) (low-pass filter at 500 Hz and high-pass filter at 5 kHz). Spontaneous activity was recorded for 3 minutes to establish baseline. Stimulation was done in three alternating 3-min epochs (OFF-ON-OFF; same parameters as those used in behavior). Data sampling was performed using a CED Micro1401 interface and Spike2 software (Cambridge Electronic Design, Cambridge, UK).

Neuronal instantaneous firing was defined as rate of the *i*-th neuron as given by *r*i (ak, bk) = *h* (ui, ak, bk, w), where *h* is a histogram function over the vector ui which stores the spiking times of the *i*-th neuron in the population, within the time interval [ak, bk), and w was the bin size for *h* (w=1s)

. Firing rate peristimulus histograms (PSTH) were calculated for the first OFF period (180s prior to stimulation) the ON period (180s of stimulation) and the second OFF period, using a bin size of 1s. In order to calculate the PSTH, each recorded spike train from a single neuron was aligned by the onset of optical stimulation. For each neuronal instantaneous firing rate *r*i the average activity during baseline was subtracted (*r*i=*r*i-avg(*r*i[*t*<180s]). Neurons were considered responsive or non-responsive to the stimulation whether their firing rate changed at least 20% from the baseline period activity (Benazzouz et al., 2000; Soares-Cunha et al., 2016).

NAc neurons were segregated into putative fast-spiking interneurons (pFSs; basal firing rate >10Hz, waveform half-width <100μs), tonically active putative cholinergic interneurons (pCINs; waveform half-width >300μs) and putative MSNs (pMSNs; basal firing rate <5Hz, that do not meet the waveform criteria for pCIN or pFS neurons) (Inokawa et al., 2010; Jin et al., 2014; Vicente et al., 2016). Given the very small representation of CINs and FSs in the recordings, only data for MSNs is shown.

VP putative GABAergic projection neurons were identified as having a baseline firing rate of 13.2±7.3Hz and action potential of 2.4±0.2ms (Pang et al., 1998). Other non-identified neurons were excluded from the analysis.

VTA neurons were segregated into putative dopaminergic (basal firing rate <10Hz, waveform duration higher than 1.5ms) and putative GABAergic (basal firing rate >10Hz, waveform duration smaller than 1.5ms) (Totah et al., 2013; Ungless, 2004). Other non-identified neurons were excluded from the analysis.

### Immunofluorescence (IF)

Mice were deeply anesthetized and then transcardially perfused with 0.9% saline followed by 4% paraformaldehyde (PFA). Brains were removed, post-fixed in 4% PFA for 24hours, transferred to 30% sucrose, for posterior sectioning in a vibratome.

Brains were sectioned coronally at a thickness of 40μm in a vibrating microtome (VT1000S, Leica, Germany). Sections containing the NAc, VP or the VTA were washed with Phosphate-Buffered Saline (PBS), treated with citrate buffer, permeabilized with PBS with 0,3% Triton X-100, blocked with PBS-T plus 10% fetal bovine serum (FBS), and then incubated with the primary antibody goat anti-GFP (1:500, ab6673; Abcam, Cambridge, UK). After washes with PBS-T, sections were incubated with the secondary fluorescent antibody Alexa Fluor^®^ 488 donkey anti-goat (1:500, A-11055, Invitrogen, Carlsbad, USA). All sections were stained with 4’,6-diamidino-2-phenylindole (1mg/ml; Invitrogen, MA, USA) and mounted using Permafluor (Invitrogen, MA, USA). Images were collected and analyzed by confocal microscopy (Olympus FluoViewTMFV3000).

Quantification of infection in each area was performed using ImageJ1.42 software [33]. For VP and VTA, percentage of area labelled with GFP was calculated by “Analyze Particles” tool, through percentage of labelled area within a given selection, obtained by defining a pixel intensity threshold for the background.

### Statistical Analysis

Prior to any statistical comparison between groups, the presence of outliers was tested (Box & Whiskers graph followed by the Tukey test). If any outlier was present, it was removed before advancing in the data analysis. The final number of animals used for statistical analysis in each figure panel is depicted in Supplementary Table 1.

After, normality tests (Shapiro-Wilks (S-W)) were performed and statistical analysis was conducted accordingly.

For histology data, comparison between two groups was done using Student’s *t*-test (when normality assumptions were not met Mann-Whitney was performed instead).

For behavioral data, Two-Way Analysis of Variance (ANOVA; Mixed Model) was used for comparisons within-between groups (OFF vs ON vs OFF; ChR2 vs YFP), and Bonferroni’s post hoc multiple comparisons was used for group differences determination. When normality assumptions were not met a mixed-effects model with the Geisser-Greenhouse correction was performed, and Bonferroni’s multiple comparison for post hoc analysis.

For comparison of neuronal populations that responded by increasing, decreasing, or not changing activity, One-Way ANOVA with repeated measures was performed, followed by Bonferroni’s *post hoc* multiple comparisons. When normality assumptions were not met, Friedman’s test was performed, followed by Dunn’s multiple comparison. For the analysis of electrophysiological temporal variation, for each time bin, the activity during stimulation was considered significant when on that time bin the activity was out 95% of the distribution of the baseline activity. Kolmogorov-Smirnov for 2 samples was performed to determine differences between the distribution of the stimulus period and the baseline.

Results are presented as mean±SEM and were considered statistically significant for *p*≤0.05. All the statistical details of experiments and final number of animals (excluding outliers) can be found throughout the results description or supplementary tables 1 and 2; these include the statistical tests used and exact *p* value.

All statistical analyses were performed using GraphPad (Prism8.0.2, La Jolla, USA).

## Acknowledgements

CS-C and BC have Scientific Employment Stimulus Contracts from the Portuguese Foundation for Science and Technology (FCT) (CEECIND/03887/2017; CEECIND/03898/2020). AVD has an FCT PhD grant (SFRH/BD/147066/2019).

This work was funded by Bial Foundation grants 30/2016 and 175/2020; by FCT under the scope of the projects PTDC/MED-NEU/29071/2017 (REWSTRESS) and PTDC/MED-NEU/4804/2020 (ENDOPIO). Part of the work has received funding from “la Caixa” Foundation (ID 100010434), under the agreement LCF/PR/HR20/52400020. This project has received funding from the European Research Council (ERC) under the European Union’s Horizon 2020 research and innovation programme (grant agreement No 101003187).

Part of the work was also funded by the ICVS Scientific Microscopy Platform, member of the national infrastructure PPBI - Portuguese Platform of Bioimaging (PPBI-POCI-01-0145-FEDER-022122); and by National funds, through the FCT - project UIDB/50026/2020 and UIDP/50026/2020.

## Competing interests

No competing interests to report.

## Author Contributions

A.J.R. and C.S.C. designed and supervised the work. R.C., C.S.C., B.C., A.V.D., and N.V.G. acquired data. R.C. analyzed, and interpreted the data. R.C., C.S.C and A.J.R. wrote the first draft of the manuscript; A.J.R., N.S., L.P. and C.S.C. revised and approved the submitted version of the manuscript. C.S.C., A.J.R. and N.S. secured funding.

## Additional files

Supplementary File 1. Statistical summary of results

---

**Figure 1 – figure supplement 1.**
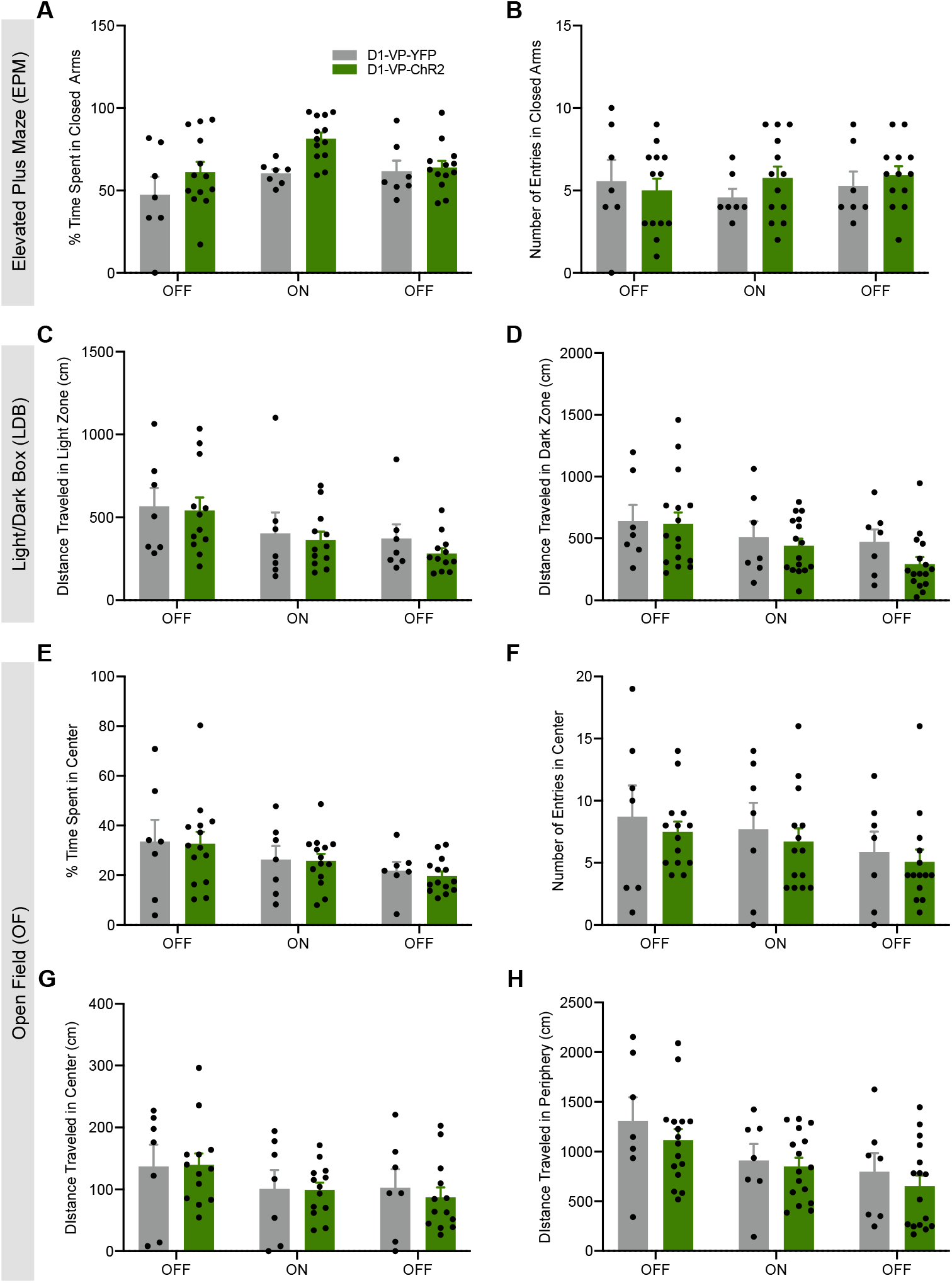
Optogenetic stimulation of D1-MSNs projecting to the VP does not change anxious-like behaviour. Optical stimulation of D1-MSN-VP projections does not alter anxiety levels in the EPM, measured by **A** % of time spent in the closed arms and **B** number of entries in the closed arms. Absence of effect of optical stimulation in the LDB test, measured by distance travelled in the **C** light zone and in the **D** dark zone of the arena. No differences were observed in the open field to what concerns **D** % of time spent and **E** number of entries in the centre of the arena, or distance travelled in the **F** centre and in the **G** periphery of the arena.Data denote mean±SEM. *p≤0.05, **p≤0.01, ***p≤0.001.

**Figure 2 – figure supplement 1.**
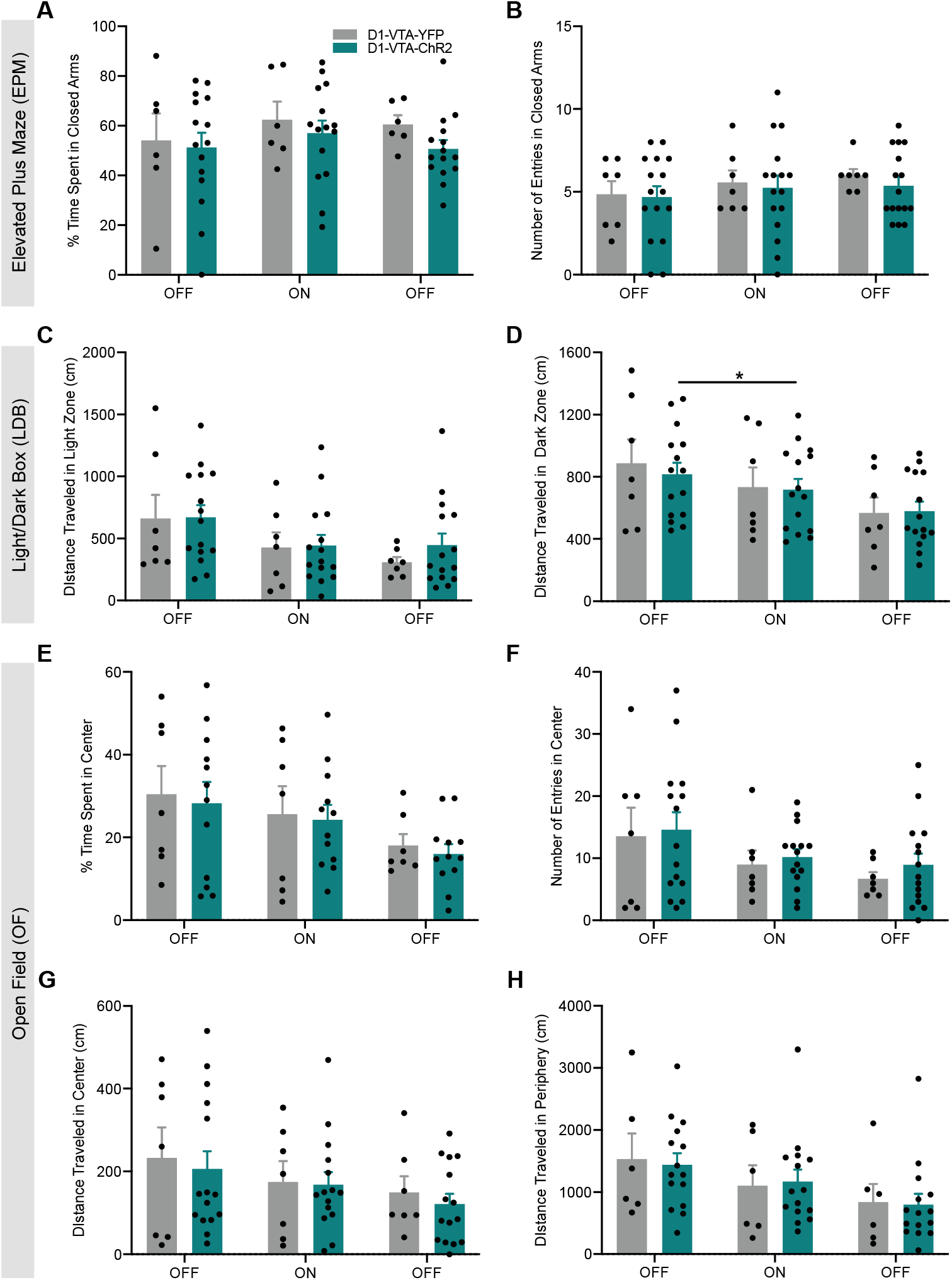
Optogenetic stimulation of D1-MSNs projecting to the VTA does not change anxious-like behaviour. Optical stimulation of D1-MSN-VTA projections does not alter anxiety levels in the EPM, measured by **A** % of time spent in the closed arms and **B** number of entries in the closed arms. Absence of effect of optical stimulation in the LDB test, measured by distance travelled in the **C** light zone and in the **D** dark zone. No differences were observed in the open field to what concerns **E** % and of time spent in the centre of the arena, **F** number of entries in the centre of the arena, **G** distance travelled in the centre of the arena and **H** distance travelled in the periphery of the arena. Data denote mean±SEM. *p≤0.05, **p≤0.01, ***p≤0.001.

**Figure 3 – figure supplement 1.**
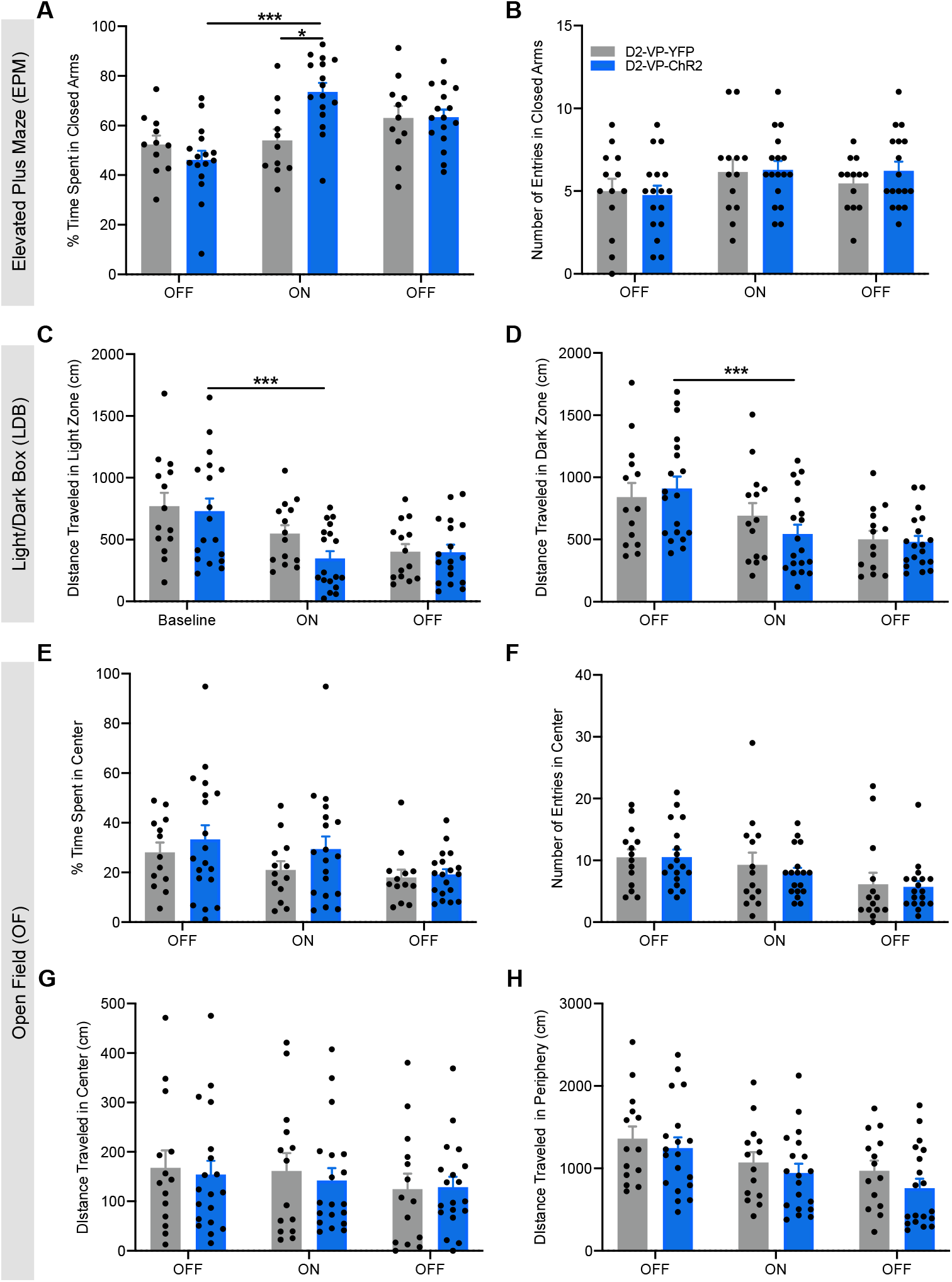
Optogenetic stimulation of D2-MSNs projecting to the VP induces an anxious-like phenotype. Significant differences were observed in terms of **A** time spent in closed arms and **B** number of entries in the closed arms of the EPM. Distance travelled in the **C** light zone and in the **D** dark zone of the LDB arena. No differences were observed in the open field to what concerns **D** % of time spent and **E** number of entries in the centre of the arena, or distance travelled in the **F** centre and in the **G** periphery of the arena. Data denote mean±SEM. *p≤0.05, ***p≤0.001.

**Figure 4 – figure supplement 1.**
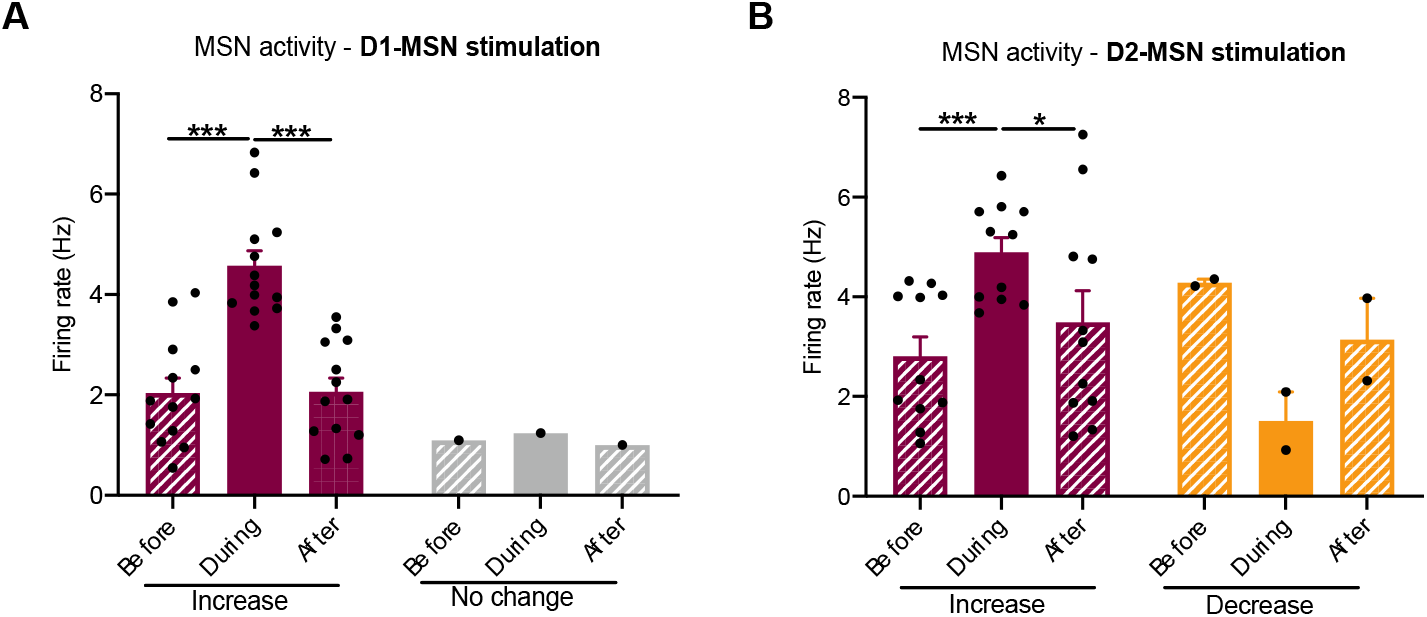
D1 - or D2-MSN optical stimulation. **A** optogenetic activation of D1-MSns significantly increased the firing rate of the majority of putative MSNs. **B** optogenetic activation of D2-MSNs significantly increased the firing rate of the majority of putative MSNs. Data denote mean±SEM. *p≤0.05, **p≤0.01, ***p≤0.001.

**Figure 7 - figure supplement 1.**
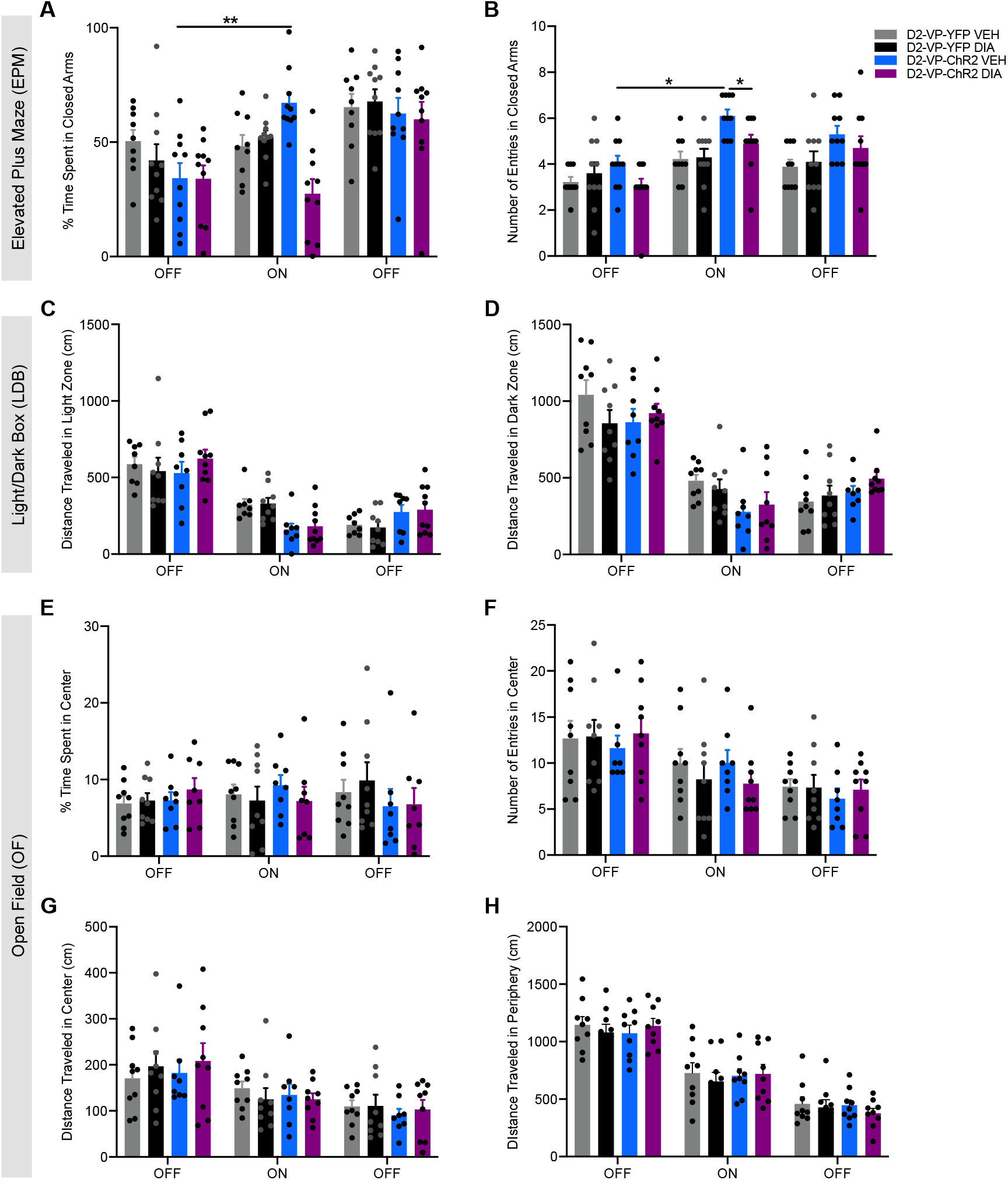
Pre-treatment with diazepam (DIA) blocks the anxious-like phenotype induced by optogenetic stimulation of D2-MSN-VP terminals. DIA treatment did not significantly alter **A** the time spent and **B** the number of entries in the closed arms of the EPM. **D-E** DIA treatment did not alter the performance in the LDB test. No differences were observed in the open field to what concerns **F** % of time spent in the centre of the arena, **G** number of entries in the centre of the arena, **H** distance travelled in the centre of the arena and **I** distance travelled in the periphery of the arena. Data denote mean±SEM. *p≤0.05, **p≤0.01, ***p≤0.001.

## Notes

### Competing Interest Statement

The authors have declared no competing interest.

## References

American Psychiatric Association. 2013. Diagnostic and Statistical Manual for Mental Disorders. 5. Washington, DC: American Psychiatric Association.

Benazzouz A, Gao DM, Ni ZG, Piallat B, Bouali-Benazzouz R, Benabid AL. 2000. Effect of high-frequency stimulation of the subthalamic nucleus on the neuronal activities of the substantia nigra pars reticulata and ventrolateral nucleus of the thalamus in the rat. Neuroscience 99:289–295. doi:10.1016/S0306-4522(00)00199-8

Berchio C, Rodrigues J, Strasser A, Michel CM, Sandi C. 2019. Trait anxiety on effort allocation to monetary incentives: a behavioral and high-density EEG study. Transl Psychiatry 9:174. doi:10.1038/s41398-019-0508-4

Bewernick BH, Kayser S, Sturm V, Schlaepfer TE. 2012. Long-Term Effects of Nucleus Accumbens Deep Brain Stimulation in Treatment-Resistant Depression: Evidence for Sustained Efficacy. Neuropsychopharmacol 37:1975–1985. doi:10.1038/npp.2012.44

Burkhouse KL, Jagan Jimmy, Defelice N, Klumpp H, Ajilore O, Hosseini B, Fitzgerald KD, Monk CS, Phan KL. 2020. Nucleus accumbens volume as a predictor of anxiety symptom improvement following CBT and SSRI treatment in two independent samples. Neuropsychopharmacol 45:561–569. doi:10.1038/s41386-019-0575-5

Cerri DH, Saddoris MP, Carelli RM. 2014. Nucleus accumbens core neurons encode value-independent associations necessary for sensory preconditioning. Behavioral Neuroscience 128:567–578. doi:10.1037/a0037797

Epstein J, Pan H, Kocsis JH, Yang Y, Butler T, Chusid J, Hochberg H, Murrough J, Strohmayer E, Stern E, Silbersweig DA. 2006. Lack of Ventral Striatal Response to Positive Stimuli in Depressed Versus Normal Subjects. AJP 163:1784–1790. doi:10.1176/ajp.2006.163.10.1784

Felix-Ortiz AC, Burgos-Robles A, Bhagat ND, Leppla CA, Tye KM. 2016. Bidirectional modulation of anxiety-related and social behaviors by amygdala projections to the medial prefrontal cortex. Neuroscience 321: 197–209. doi:10.1016/j.neuroscience.2015.07.041

Feng J, Pena CJ, Purushothaman I, Engmann O, Walker D, Brown AN, Issler O, Doyle M, Harrigan E, Mouzon E, Vialou V, Shen L, Dawlaty MM, Jaenisch R, Nestler EJ. 2017. Tet1 in Nucleus Accumbens Opposes Depression-and Anxiety-Like Behaviors. Neuropsychopharmacol 42:1657–1669. doi:10.1038/npp.2017.6

Francis TC, Chandra R, Friend DM, Finkel E, Dayrit G, Miranda J, Brooks JM, Iñiguez SD, O’Donnell P, Kravitz A, Lobo MK. 2015. Nucleus Accumbens Medium Spiny Neuron Subtypes Mediate Depression-Related Outcomes to Social Defeat Stress. Biological Psychiatry 77:212–222. doi:10.1016/j.biopsych.2014.07.021

Hein TP, de Fockert J, Ruiz MH. 2021. State anxiety biases estimates of uncertainty and impairs reward learning in volatile environments. NeuroImage 224:117424. doi:10.1016/j.neuroimage.2020.117424

Heshmati M, Golden SA, Pfau ML, Christoffel DJ, Seeley EL, Cahill ME, Khibnik LA, Russo SJ. 2016. Mefloquine in the nucleus accumbens promotes social avoidance and anxiety-like behavior in mice. Neuropharmacology 101:351–357. doi: 10.1016/j.neuropharm.2015.10.013

Inokawa H, Yamada H, Matsumoto N, Muranishi M, Kimura M. 2010. Juxtacellular labeling of tonically active neurons and phasically active neurons in the rat striatum. Neuroscience 168:395–404. doi:10.1016/j.neuroscience.2010.03.062

Jin X, Tecuapetla F, Costa RM. 2014. Basal ganglia subcircuits distinctively encode the parsing and concatenation of action sequences. Nat Neurosci 17:423–430. doi:10.1038/nn.3632

Kilts CD, Kelsey JE, Knight B, Ely TD, Bowman FD, Gross RE, Selvig A, Gordon A, Newport DJ, Nemeroff CB. 2006. The Neural Correlates of Social Anxiety Disorder and Response to Pharmacotherapy. Neuropsychopharmacol 31:2243–2253. doi:10.1038/sj.npp.1301053

Kim S-Y, Adhikari A, Lee SY, Marshel JH, Kim CK, Mallory CS, Lo M, Pak S, Mattis J, Lim BK, Malenka RC, Warden MR, Neve R, Tye KM, Deisseroth K. 2013. Diverging neural pathways assemble a behavioural state from separable features in anxiety. Nature 496:219–223. doi:10.1038/nature12018

Kühn S, Schubert F, Gallinat J. 2011. Structural correlates of trait anxiety: Reduced thickness in medial orbitofrontal cortex accompanied by volume increase in nucleus accumbens. Journal of Affective Disorders 134:315–319. doi:10.1016/j.jad.2011.06.003

Kupchik YM, Brown RM, Heinsbroek JA, Lobo MK, Schwartz DJ, Kalivas PW. 2015. Coding the direct/indirect pathways by D1 and D2 receptors is not valid for accumbens projections. Nat Neurosci 18:1230–1232. doi:10.1038/nn.4068

Levita L, Hoskin R, Champi S. 2012. Avoidance of harm and anxiety: A role for the nucleus accumbens. NeuroImage 62:189–198. doi:10.1016/j.neuroimage.2012.04.059

Li Z, Chen Z, Fan G, Li A, Yuan J, Xu T. 2018. Cell-Type-Specific Afferent Innervation of the Nucleus Accumbens Core and Shell. Front Neuroanat 12:84. doi:10.3389/fnana.2018.00084

Lim BK, Huang KW, Grueter BA, Rothwell PE, Malenka RC. 2012. Anhedonia requires MC4R-mediated synaptic adaptations in nucleus accumbens. Nature 487:183–189. doi:10.1038/nature11160

Löw K, Crestani F, Keist R, Benke D, Brünig I, Benson JA, Fritschy J-M, Rülicke T, Bluethmann H, Möhler H, Rudolph U. 2000. Molecular and Neuronal Substrate for the Selective Attenuation of Anxiety. Science 290:131–134. doi:10.1126/science.290.5489.131

Moaddab M, Ray MH, McDannald MA. 2021. Ventral pallidum neurons dynamically signal relative threat. Commun Biol 4:43. doi:10.1038/s42003-020-01554-4

Morrison SE, McGinty VB, du Hoffmann J, Nicola SM. 2017. Limbic-motor integration by neural excitations and inhibitions in the nucleus accumbens. Journal of Neurophysiology 118:2549–2567. doi:10.1152/jn.00465.2017

O’Brien CP, Childress AR, Ehrman R, Robbins SJ. 1998. Conditioning factors in drug abuse: can they explain compulsion? J Psychopharmacol 12:15–22. doi:10.1177/026988119801200103

Pádua-Reis M, Nôga DA, Tort ABL, Blunder M. 2021. Diazepam causes sedative rather than anxiolytic effects in C57BL/6J mice. Sci Rep 11:9335. doi:10.1038/s41598-021-88599-5

Pang K, Tepper JM, Zaborszky L. 1998. Morphological and electrophysiological characteristics of noncholinergic basal forebrain neurons. J Comp Neurol 394:186–204. doi:10.1002/(SICI)1096-9861(19980504)394:2<186::AID-CNE4>3.0.CO;2-Z

Paxinos G, Franklin K. n.d. The Mouse Brain in Stereotaxic Coordinates, 3rd Edition. ed. Academic Press.

Qi G, Zhang P, Li T, Li M, Zhang Q, He F, Zhang L, Cai H, Lv X, Qiao H, Chen X, Ming J, Tian B. 2022. NAc-VTA circuit underlies emotional stress-induced anxiety-like behavior in the three-chamber vicarious social defeat stress mouse model. Nat Commun 13:577. doi:10.1038/s41467-022-28190-2

Russo SJ, Nestler EJ. 2013. The brain reward circuitry in mood disorders. Nat Rev Neurosci 14:609–625. doi:10.1038/nrn3381

Salamone JD, Correa M. 2012. The Mysterious Motivational Functions of Mesolimbic Dopamine. Neuron 76:470–485. doi:10.1016/j.neuron.2012.10.021

Satterthwaite TD, Kable JW, Vandekar L, Katchmar N, Bassett DS, Baldassano CF, Ruparel K, Elliott MA, Sheline YI, Gur RC, Gur RE, Davatzikos C, Leibenluft E, Thase ME, Wolf DH. 2015. Common and Dissociable Dysfunction of the Reward System in Bipolar and Unipolar Depression. Neuropsychopharmacol 40:2258–2268. doi:10.1038/npp.2015.75

Shen EY, Jiang Y, Javidfar B, Kassim B, Loh Y-HE, Ma Q, Mitchell AC, Pothula V, Stewart AF, Ernst P, Yao W-D, Martin G, Shen L, Jakovcevski M, Akbarian S. 2016. Neuronal Deletion of Kmt2a/Mll1 Histone Methyltransferase in Ventral Striatum is Associated with Defective Spike-Timing-Dependent Striatal Synaptic Plasticity, Altered Response to Dopaminergic Drugs, and Increased Anxiety. Neuropsychopharmacol 41:3103–3113. doi:10.1038/npp.2016.144

Soares-Cunha C, Coimbra B, David-Pereira A, Borges S, Pinto L, Costa P, Sousa N, Rodrigues AJ. 2016. Activation of D2 dopamine receptor-expressing neurons in the nucleus accumbens increases motivation. Nat Commun 7:11829. doi:10.1038/ncomms11829

Soares-Cunha C, Coimbra B, Domingues AV, Vasconcelos N, Sousa N, Rodrigues AJ. 2018. Nucleus Accumbens Microcircuit Underlying D2-MSN-Driven Increase in Motivation. eNeuro 5:ENEURO.0386-18.2018. doi:10.1523/ENEURO.0386-18.2018

Soares-Cunha C, de Vasconcelos NAP, Coimbra B, Domingues AV, Silva JM, Loureiro-Campos E, Gaspar R, Sotiropoulos I, Sousa N, Rodrigues AJ. 2020. Nucleus accumbens medium spiny neurons subtypes signal both reward and aversion. Mol Psychiatry 25:3241–3255. doi:10.1038/s41380-019-0484-3

Soares-Cunha C, Domingues AV, Correia R, Coimbra B, Vieitas-Gaspar N, de Vasconcelos NAP, Pinto L, Sousa N, Rodrigues AJ. 2022. Distinct role of nucleus accumbens D2-MSN projections to ventral pallidum in different phases of motivated behavior. Cell Reports 38:110380. doi:10.1016/j.celrep.2022.110380

Tan KR, Yvon C, Turiault M, Mirzabekov JJ, Doehner J, Labouèbe G, Deisseroth K, Tye KM, Lüscher C. 2012. GABA Neurons of the VTA Drive Conditioned Place Aversion. Neuron 73:1173–1183. doi:10.1016/j.neuron.2012.02.015

Totah NKB, Kim Y, Moghaddam B. 2013. Distinct prestimulus and poststimulus activation of VTA neurons correlates with stimulus detection. Journal of Neurophysiology 110:75–85. doi:10.1152/jn.00784.2012

Tye KM, Prakash R, Kim S-Y, Fenno LE, Grosenick L, Zarabi H, Thompson KR, Gradinaru V, Ramakrishnan C, Deisseroth K. 2011. Amygdala circuitry mediating reversible and bidirectional control of anxiety. Nature 471:358–362. doi:10.1038/nature09820

Ungless MA. 2004. Uniform Inhibition of Dopamine Neurons in the Ventral Tegmental Area by Aversive Stimuli. Science 303:2040–2042. doi:10.1126/science.1093360

Vicente AM, Galvão-Ferreira P, Tecuapetla F, Costa RM. 2016. Direct and indirect dorsolateral striatum pathways reinforce different action strategies. Current Biology 26:R267–R269. doi:10.1016/j.cub.2016.02.036

West EA, Carelli RM. 2016. Nucleus Accumbens Core and Shell Differentially Encode Reward-Associated Cues after Reinforcer Devaluation. Journal of Neuroscience 36:1128–1139. doi:10.1523/JNEUROSCI.2976-15.2016

Wulff AB, Tooley J, Marconi LJ, Creed MC. 2019. Ventral pallidal modulation of aversion processing. Brain Research 1713:62–69. doi:10.1016/j.brainres.2018.10.010

Zweifel LS, Fadok JP, Argilli E, Garelick MG, Jones GL, Dickerson TMK, Allen JM, Mizumori SJY, Bonci A, Palmiter RD. 2011. Activation of dopamine neurons is critical for aversive conditioning and prevention of generalized anxiety. Nat Neurosci 14:620–626. doi:10.1038/nn.2808

