## Supplementary Table for "Involvement of nucleus accumbens D2-MSN projections to the ventral pallidum in anxious-like behavior"

### Supplementary File 1

#### **Table 1.** Statistical summary of main figures

| Figure | Number of animals | Statistical test | Test Value | *Post hoc* |
|---|---|---|---|---|
| Figure 1C | D1-VP-ChR2=8<br>D1-VP-YFP=5 | Unpaired t-test | $t_{11}$=0.8, p=0.4390 | |
| Figure 1E | D1-VP-ChR2=12<br>D1-VP-YFP=7 | Two-way ANOVA | Stim: $F_{2,22}$=8.8, p=0.0016<br>Group: $F_{1,11}$=0.1, p=0.7184 | OFF ChR2 vs ON ChR2 p=0.0020 |
| Figure 1F | D1-VP-ChR2=14<br>D1-VP-YFP=7 | Two-way ANOVA | Stim: $F_{2,26}$=0.4, p=0.0007<br>Group: $F_{1,13}$=3.1, p=6878 | |
| Figure 1H | D1-VP-ChR2=14<br>D1-VP-YFP=7 | Two-way ANOVA | Stim: $F_{2,30}$=37.8, p<0.0001<br>Group: $F_{1,13}$=3.1, p=0.1490 | OFF ChR2 vs ON ChR2, p=0.0018<br>OFF ChR2 vs OFF(2) ChR2, p<0.0001<br>OFF ChR2 vs OFF(2) YFP, p=0.0001<br>OFF YFP vs OFF(2) YFP, p=0.0004<br>ON ChR2 vs OFF(2) ChR2, p=0.0059<br>ON ChR2 vs OFF(2) YFP, p=0.0486<br>ON YFP vs OFF(2), p=0.0336 |
| Figure 1I | D1-VP-ChR2=14<br>D1-VP-YFP=7 | Two-way ANOVA | Stim: $F_{2,26}$=20.0, p<0.0001<br>Group: $F_{1,13}$=3.1, p=0.1028 | OFF ChR2 vs ON ChR2, p=0.0006<br>OFF ChR2 vs ON YFP, p=0.0009<br>OFF ChR2 vs OFF(2) ChR2, p<0.0001<br>OFF ChR2 vs OFF(2) YFP, p<0.0001<br>OFF YFP vs OFF(2) YFP, p=0.0283 |
| Figure 1K | D1-VP-ChR2=16<br>D1-VP-YFP=7 | Two-way ANOVA | Stim: $F_{1,22}$=0.1, p=0.7038<br>Group: $F_{1,22}$=0.1, p=0.8067 | |
| Figure 1L | D1-VP-ChR2=15<br>D1-VP-YFP=7 | Two-way ANOVA | Stim: $F_{1,20}$=0.2, p=0.6260<br>Group: $F_{1,20}$=0.0, p=0.8519 | |
| Figure 1M | D1-VP-ChR2=15<br>D1-VP-YFP=7 | Two-way ANOVA | Stim: $F_{1,20}$=2.1, p=0.1611<br>Group: $F_{1,20}$=1.4, p=0.2574 | |
| Figure 2C | D1-VTA-ChR2=6<br>D1-VTA-YFP=5 | Unpaired t-test | $t_{9}$=0.4, p=0.6991 | |
| Figure 2E | D1-VTA-ChR2=14<br>D1-VTA-YFP=7 | Two-way ANOVA | Stim: $F_{1,26}$=0.4, p=0.6845<br>Group: $F_{1,13}$=3.1, p=0.1019 | |
| Figure 2F | D1-VTA-ChR2=14<br>D1-VTA-YFP=7 | Two-way ANOVA | Stim: $F_{1,26}$=2.2, p=0.1275<br>Group: $F_{1,13}$=0.7, p=0.4065 | |
| Figure 2H | D1-VTA-ChR2=13<br>D1-VTA-YFP=7 | Two-way ANOVA | Stim: $F_{1,24}$=46.6, p<0.0001<br>Group: $F_{1,12}$=0.2, p=0.6914 | OFF ChR2 vs OFF(2) ChR2, p<0.0001<br>OFF ChR2 vs OFF(2) YFP, p=0.0026<br>OFF YFP vs OFF(2) ChR2, p=0.0148<br>OFF YFP vs OFF(2) YFP, p<0.0001<br>ON ChR2 vs OFF(2) ChR2, p=0.0017<br>ON YFP vs OFF(2) YFP, p=0.0132 |
| Figure 2I | D1-VTA-ChR2=15<br>D1-VTA-YFP=7 | Two-way ANOVA | Stim: $F_{2,28}$=38.4, p<0.0001<br>Group: $F_{1,14}$=1.4, p=0.2589 | OFF ChR2 vs ON ChR2, p=0.0007<br>OFF ChR2 vs ON YFP, p=0.0153<br>OFF ChR2 vs OFF(2) ChR2, p<0.0001<br>OFF ChR2 vs OFF(2) YFP, p<0.0001<br>OFF YFP vs ON YFP, p=0.0249<br>OFF YFP vs OFF(2) ChR2, p=0.0140<br>OFF YFP vs OFF(2) YFP, p<0.0001 |
| Figure 2K | D1-VTA-ChR2=13<br>D1-VTA-YFP=7 | Two-way ANOVA | Stim: $F_{1,24}$=0.1, p=0.7761<br>Group: $F_{1,12}$=0.5, p=0.4885 | |
| Figure 2L | D1-VTA-ChR2=13<br>D1-VTA-YFP=7 | Two-way ANOVA | Stim: $F_{1,24}$=0.2, p=0.7008<br>Group: $F_{1,12}$=0.1, p=0.7402 | |
| Figure 2M | D1-VTA-ChR2=14<br>D1-VTA-YFP=7 | Two-way ANOVA | Stim: $F_{1,24}$=1.1, p=0.2872<br>Group: $F_{1,12}$=0.1, p=0.7887 | |
| Figure 3C | D2-VP-ChR2=8<br>D2-VP-YFP=5 | Unpaired t-test | $t_{11}$=0.3, p=0.7448 | |
| Figure 3E | D2-VP-ChR2=17<br>D2-VP-YFP=12 | Two-way ANOVA | Stim: $F2_{,32}$=120.3, p<0.0001<br>Group: $F_{1,16}$=1.7, p=0.2071 | OFF ChR2 vs ON ChR2, p<0.0001<br>OFF ChR2 vs ON YFP, p=0.0379<br>OFF ChR2 vs OFF(2) ChR2, p<0.0001<br>OFF ChR2 vs OFF(2) YFP, p<0.0001<br>OFF YFP vs ON ChR2, p<0.0001<br>OFF YFP vs ON YFP, p=0.0061<br>OFF YFP vs OFF(2) ChR2, p<0.0001<br>OFF YFP vs OFF(2) YFP, p<0.0001 |

| | | | | |
|---|---|---|---|---|
| | | | | ON ChR2 vs ON YFP, p=0.0002<br>ON YFP vs OFF(2) ChR2, p=0.0009<br>ON YFP vs OFF(2) YFP, p<0.0001 |
| Figure 3F | D2-VP-ChR2=16<br>D2-VP-YFP=11 | Two-way ANOVA | Stim: $F_{2,34}$=30.5, p<0.0001<br>Group: $F_{1,17}$=1.8, p=0.2028 | OFF ChR2 vs ON ChR2, p<0.0001<br>OFF ChR2 vs ON YFP, p=0.0390<br>OFF ChR2 vs OFF(2) ChR2, p<0.0001<br>OFF ChR2 vs OFF(2) YFP, p<0.0001<br>OFF YFP vs ON ChR2, p=0.0004<br>OFF YFP vs OFF(2) ChR2, p=0.004<br>OFF YFP vs OFF(2) YFP, p=0.0314<br>ON YFP vs OFF(2) ChR2, p=0.0325 |
| Figure 3H | D2-VP-ChR2=17<br>D2-VP-YFP=12 | Two-way ANOVA | Stim: $F_{2,32}$=120.3, p<0.0001<br>Group: $F_{1,16}$=1.7, p=0.2071 | OFF ChR2 vs ON ChR2, p<0.001<br>OFF ChR2 vs ON YFP, p=0.0379<br>OFF ChR2 vs OFF(2) ChR2, p<0.0001<br>OFF ChR2 vs OFF(2) YFP, p<0.0001<br>OFF ChR2 vs ON ChR2, p<0.0001<br>OFF YFP vs ON YFP, p=0.0061<br>OFF YFP vs OFF(2) ChR2, p<0.0001<br>OFF YFP vs OFF(2) YFP, p<0.0001<br>ON ChR2 vs ON YFP, p=0.0002<br>ON YFP vs OFF(2) ChR2, p=0.0009<br>ON YFP vs OFF(2) YFP, p<0.0001 |
| Figure 3I | D2-VP-ChR2=18<br>D2-VP-YFP=11 | Two-way ANOVA | Stim: $F_{2,36}$=26.9, p<0.0001<br>Group: $F_{1,18}$=0.5, p=0.4922 | OFF ChR2 vs ON ChR2, p<0.0001<br>OFF ChR2 vs OFF(2) ChR2, p<0.0001<br>OFF ChR2 vs OFF(2) YFP, p=0.0003<br>OFF YFP vs ON ChR2, p=0.0011<br>OFF YFP vs OFF(2) ChR2, p=0.0054<br>OFF YFP vs OFF(2) YFP, p=0.0002 |
| Figure 3K | D2-VP-ChR2=18<br>D2-VP-YFP=12 | Two-way ANOVA | Stim: $F_{1,28}$=13.9, p=0.0009<br>Group: $F_{1,28}$=4.5, p=0.0439 | ChR2 ON vs ChR2 OFF, p<0.0001<br>ChR2 ON vs YFP ON, p<0.0001 |
| Figure 3L | D2-VP-ChR2=15<br>D2-VP-YFP=13 | Two-way ANOVA | Stim: $F_{1,26}$=8.0, p=0.0089<br>Group: $F_{1,26}$=2.7, p=0.1123 | ChR2 ON vs ChR2 OFF, p=0.0016<br>ChR2 ON vs YFP ON, p=0.0167 |
| Figure 3M | D2-VP-ChR2=17<br>D2-VP-YFP=13 | Two-way ANOVA | Stim: $F_{1,58}$=0.3, p=0.5583<br>Group: $F_{1,58}$=0.0, p=0.9790 | |
| Figure 4D | Increase: 1 neuron<br>Decrease: 13 neurons<br>No Change: 13 neurons | One way ANOVA | Increase: $F_{2,30}$=35.8, p<0.0001 | Before vs during, p<0.0001<br>During vs after, p<0.0001 |
| Figure 5D | Increase: 7 neurons<br>Decrease: 23 neurons<br>No Change: 8 neurons | One way ANOVA | Increase: $F_{2,12}$=10.6, p=0.0092<br><br>Decrease: $F_{2,44}$=38.6, p<0.0001 | Before vs during, p=0.0449<br>During vs after, p=0.0406<br><br>Before vs during, p<0.0001<br>During vs after, p<0.0001 |
| Figure 5F | Increase: 1 neuron<br>Decrease: 3 neurons<br>No Change: 1 neuron | One way ANOVA | Decrease: $F_{2,4}$=6.7, p=0.0926 | |
| Figure 6D | Decrease: 25 neurons<br>No Change: 10 neurons | One way ANOVA | Decrease: $F_{2,24}$=16.7, p<0.0001 | Before vs during, p=0.0001<br>During vs after, p=0.0060 |
| Figure 7B | D2-VP-ChR2-VEH=10<br>D2-VP-ChR2-DIA=9<br>D2-VP-YFP VEH=9<br>D2-VP-YFP DIA=10 | Two-way ANOVA | Stim: $F_{2,18}$=29.5, p<0.0001<br>Group: $F3_{,27}$=2.1, p=0.1279 | OFF ChR2 VEH vs. ON ChR2 VEH, p=0.0002<br>OFF ChR2 VEH vs. OFF(2) ChR2 VEH, p=0.0016<br>OFF ChR2 VEH vs. OFF(2) YFP VEH=0.0186<br>OFF ChR2 VEH vs. OFF(2) ChR2 DIA, p=0.0028<br>OFF ChR2 VEH vs. OFF(2) YFP DIA, p=0.002<br>OFF ChR2 DIA vs. ON ChR2 VEH, p=0.0046<br>OFF ChR2 DIA vs. OFF(2) ChR2 VEH, p=0.025<br>OFF ChR2 DIA vs. OFF(2) YFP VEH, p=0.0328<br>OFF ChR2 DIA vs. OFF(2) ChR2 DIA, p=0.0005<br>OFF ChR2 DIA vs. OFF(2) YFP DIA, p=0.004<br>OFF YFP DIA vs. OFF(2) YFP DIA, p=0.0207<br>ON ChR2 VEH vs. ON ChR2 DIA, p<0.0001<br>ON ChR2 DIA vs. OFF(2) ChR2 VEH, p=0.0002 |

| | | | | |
|---|---|---|---|---|
| | | | | ON ChR2 DIA vs. OFF(2) YFP VEH, p=0.0002<br>ON ChR2 DIA vs. OFF(2) ChR2 DIA, p<0.0001<br>ON ChR2 DIA vs. OFF(2)YFP DIA, p<0.0001 |
| Figure 7C | D2-VP-ChR2-VEH=10<br>D2-VP-ChR2-DIA=10<br>D2-VP-YFP VEH=9<br>D2-VP-YFP DIA=10 | Two-way ANOVA | Stim: $F_{2,18}$=48.0, p<0.0001<br>Group: $F_{3,27}$=1.5, p=0.2252 | OFF YFP DIA vs. ON ChR2 VEH, p<0.0001<br>OFF YFP DIA vs. OFF(2) ChR2 VEH, p=0.0002<br>OFF YFP DIA vs. OFF(2) YFP VEH, p=0.0067<br>OFF YFP DIA vs. OFF(2) YFP DIA, p=0.0079<br>ON ChR2 VEH vs. ON YFP VEH, p=0.0332<br>ON ChR2 VEH vs. ON ChR2 DIA, p=0.0329<br>ON ChR2 VEH vs. ON YFP DIA, p=0.0079<br>ON YFP DIA vs. OFF(2) ChR2 VEH, p=0.0206 |
| Figure 7D | D2-VP-ChR2-VEH=8<br>D2-VP-ChR2-DIA=10<br>D2-VP-YFP VEH=9<br>D2-VP-YFP DIA=9 | Two-way ANOVA | Stim: $F_{2,18}$=88.1, p<0.0001<br>Group: $F_{3,27}$=1.1, p=0.0717 | OFF ChR2 VEH vs. ON ChR2 VEH. P<0.0001<br>OFF ChR2 VEH vs. ON ChR2 DIA, p=0.0021<br>OFF ChR2 VEH vs. ON YFP DIA, p=0.0097<br>OFF ChR2 VEH vs. OFF(2) ChR2 VEH, p<0.00[illegible]<br>OFF ChR2 VEH vs. OFF(2) YFP VEH, p=0.001[illegible]<br>OFF ChR2 VEH vs. OFF(2) ChR2 DIA, p=0.010[illegible]<br>OFF ChR2 VEH vs. OFF(2) YFP DIA, p=0.0003<br>OFF YFP VEH vs. ON ChR2 VEH, p<0.0001<br>OFF YFP VEH vs. ON YFP VEH, p=0.0006<br>OFF YFP VEH vs. ON ChR2 DIA, p=0.0002<br>OFF YFP VEH vs. ON YFP DIA, p=0.0012<br>OFF YFP VEH vs. OFF(2) ChR2 VEH, p=0.000[illegible]<br>OFF YFP VEH vs. OFF(2) YFP VEH, p<0.0001<br>OFF YFP VEH vs. OFF(2) ChR2 DIA, p=0.0012<br>OFF YFP VEH vs. OFF(2) YFP DIA, p<0.0001<br>OFF ChR2 DIA vs. ON ChR2 VEH, p<0.0001<br>OFF ChR2 DIA vs. ON YFP VEH, p=0.0009<br>OFF ChR2 DIA vs. ON ChR2 DIA, p<0.0001<br>OFF ChR2 DIA vs. ON YFP DIA, <0.0001<br>OFF ChR2 DIA vs. OFF(2) ChR2 VEH, <0.0001<br>OFF ChR2 DIA vs. OFF(2) YFP VEH, <0.0001<br>OFF ChR2 DIA vs. OFF(2) ChR2 DIA, <0.0001<br>OFF ChR2 DIA vs. OFF(2) YFP DIA, <0.0001<br>OFF YFP DIA vs. ON ChR2 VEH, <0.0001<br>OFF YFP DIA vs. ON YFP VEH, p=0.0239<br>OFF YFP DIA vs. ON ChR2 DIA, p=0.0003<br>OFF YFP DIA vs. ON YFP DIA, p<0.0001<br>OFF YFP DIA vs. OFF(2) ChR2 VEH, p=0.0005<br>OFF YFP DIA vs. OFF(2) YFP VEH, p=0.0002<br>OFF YFP DIA vs. OFF(2) ChR2 DIA, p=0.0016<br>OFF YFP DIA vs. OFF(2) YFP DIA, p<0.0001 |
| Figure 7E | D2-VP-ChR2-VEH=8<br>D2-VP-ChR2-DIA=10<br>D2-VP-YFP VEH=8<br>D2-VP-YFP DIA=9 | Two-way ANOVA | Stim: $F_{2,18}$=67.5, p<0.0001<br>Group: $F_{3,27}$=0.2, p=0.8953 | OFF ChR2 VEH vs. ON ChR2 VEH, p=0.0112<br>OFF ChR2 VEH vs. OFF(2) YFP DIA, p=0.0491<br>OFF YFP VEH vs. ON YFP VEH, p<0.0001<br>OFF YFP VEH vs. ON ChR2 DIA, p=0.0221<br>OFF YFP VEH vs. OFF(2) YFP VEH, p=0.0001<br>OFF YFP VEH vs. OFF(2) YFP DIA, p=0.0054<br>OFF ChR2 DIA vs. ON ChR2 VEH, p=0.03<br>OFF ChR2 DIA vs. ON YFP VEH, p=0.0054<br>OFF ChR2 DIA vs. ON ChR2 DIA, p<0.0001<br>OFF ChR2 DIA vs. ON YFP DIA, p=0.0215<br>OFF ChR2 DIA vs. OFF(2) YFP VEH, p=0.0124<br>OFF ChR2 DIA vs. OFF(2) ChR2 DIA, p=0.0002<br>OFF ChR2 DIA vs. OFF(2) YFP DIA, p=0.0006<br>OFF YFP DIA vs. ON YFP DIA, p=0.0022<br>OFF YFP DIA vs. OFF YFP DIA, p<0.0001 |
| Figure 7F | D2-VP-ChR2-VEH=10 | Two-way ANOVA | Stim: $F_{1,9}$=49.05, p<0.0001<br>Group: $F_{3,27}$=35.36, p<0.0001 | OFF ChR2 VEH vs. ON ChR2 VEH, p<0.000<br>OFF ChR2 DIA vs. ON ChR2 VEH, p<0.0001 |

| | | | | |
|---|---|---|---|---|
| | D2-VP-ChR2-DIA=110<br>D2-VP-YFP VEH=10<br>D2-VP-YFP DIA=9 | | | OFF YFP DIA vs. ON ChR2 VEH, $p<0.0001$<br>OFF YFP VEH vs. ON ChR2 VEH, $p<0.0001$<br>ON ChR2 VEH vs. ON ChR2 DIA, $p<0.0001$<br>ON ChR2 VEH vs. ON YFP DIA, $p<0.0001$<br>ON ChR2 VEH vs. ON YFP VEH, $p<0.0001$ |
| Figure 7G | D2-VP-ChR2-VEH=10<br>D2-VP-ChR2-DIA=110<br>D2-VP-YFP VEH=10<br>D2-VP-YFP DIA=9 | Two-way ANOVA | Stim: $F_{1,9}=7.1$, $p=0.0253$<br>Group: $F_{3,27}=9.0$, $p=0.0003$ | OFF ChR2 VEH vs. ON ChR2 VEH, $p=0.0003$<br>OFF ChR2 DIA vs. ON ChR2 VEH, $p<0.0001$<br>OFF YFP DIA vs. ON ChR2 VEH, $p=0.0001$<br>OFF YFP VEH vs. ON ChR2 VEH, $p<0.0001$<br>ON ChR2 VEH vs. ON ChR2 DIA, $p<0.0001$<br>ON ChR2 VEH vs. ON YFP DIA, $p=0.001$<br>ON ChR2 VEH vs. ON YFP VEH, $p<0.0001$ |
| Figure 7H | D2-VP-ChR2-VEH=9<br>D2-VP-ChR2-DIA=8<br>D2-VP-YFP VEH=9<br>D2-VP-YFP DIA=7 | Two-way ANOVA | Stim: $F_{1,9}=0.001$, $p=0.9729$<br>Group: $F_{3,27}=3.0$, $p=0.051$ | |

**Table 2.** Statistical summary of figure supplements

| Figure | Number of animals | Statistical test | Test Value | *Post hoc* |
|---|---|---|---|---|
| Figure 1 – figure supplement 1 A | D1-VP-ChR2=13<br>D1-VP-YFP=7 | Two-way ANOVA | Stim: $F_{2,24}$=4.5, p=0.0213<br>Group: $F_{1,12}$=9.3, p=0.0101 | OFF YFP vs. ON ChR2, p=0.0007 |
| Figure 1 – figure supplement 1 B | D1-VP-ChR2=13<br>D1-VP-YFP=7 | Two-way ANOVA | Stim: $F_{2,24}$=0.2, p=0.8158<br>Group: $F_{1,12}$=0.3, p=0.603 | |
| Figure 1 – figure supplement 1 C | D1-VP-ChR2=12<br>D1-VP-YFP=7 | Two-way ANOVA | Stim: $F_{2,24}$=8.2, p=0.0019<br>Group: $F_{1,12}$=0.1711, p=0.6864 | OFF ChR2 vs. OFF ChR2, p=0.0078 |
| Figure 1 – figure supplement 1 D | D1-VP-ChR2=16<br>D1-VP-YFP=7 | Two-way ANOVA | Stim: $F_{2,30}$=8.6, p=0.0011<br>Group: $F_{1,15}$=0.4, p=0.541 | OFF ChR2 vs. OFF ChR2, p=0.0004 |
| Figure 1 – figure supplement 1 E | D1-VP-ChR2=14<br>D1-VP-YFP=7 | Two-way ANOVA | Stim: $F_{2,26}$=6.3, p=0.0059<br>Group: $F_{1,13}$=0.05, p=0.8210 | OFF ChR2 vs. OFF ChR2, p=0.0454 |
| Figure 1 – figure supplement 1 F | D1-VP-ChR2=14<br>D1-VP-YFP=7 | Two-way ANOVA | Stim: $F_{2,26}$=4.2, p=0.0266<br>Group: $F_{1,13}$=0.3, p=0.5787 | |
| Figure 1 – figure supplement 1 G | D1-VP-ChR2=13<br>D1-VP-YFP=7 | Two-way ANOVA | Stim: $F_{2,24}$=6.7, p=0.0048<br>Group: $F_{1,12}$=0.1, p=0.9212 | OFF ChR2 vs. OFF ChR2, p=0.0272 |
| Figure 1 – figure supplement 1 H | D1-VP-ChR2=16<br>D1-VP-YFP=7 | Two-way ANOVA | Stim: $F_{2,30}$=17.8, p<0.0001<br>Group: $F_{1,15}$=0.6, p=0.4549 | OFF ChR2 vs. OFF(2) ChR2, p=0.0002<br>OFF ChR2 vs. OFF(2) YFP, p=0.0113<br>OFF YFP vs. OFF(2) YFP, p=0.0092 |
| Figure 2 – figure supplement 1 A | D1-VTA-ChR2=15<br>D1-VTA-YFP=7 | Two-way ANOVA | Stim: $F_{2,28}$=1.0, p=0.3785<br>Group: $F_{1,14}$=0.7, p=0.4164 | OFF YFP vs. ON ChR2 p=0.0007 |
| Figure 2 – figure supplement 1 B | D1-VTA-ChR2=16<br>D1-VTA-YFP=7 | Two-way ANOVA | Stim: $F_{2,30}$=1.8, p=0.1874<br>Group: $F_{1,15}$=1.0, p=0.3253 | |
| Figure 2 – figure supplement 1 C | D1-VTA-ChR2=15<br>D1-VTA-YFP=7 | Two-way ANOVA | Stim: $F_{2,28}$=4.2, p=0.8158<br>Group: $F_{1,14}$=0.2, p=0.6986 | |
| Figure 2 – figure supplement 1 D | D1-VTA-ChR2=15<br>D1-VTA-YFP=7 | Two-way ANOVA | Stim: $F_{2,28}$=62.22, p<0.0001<br>Group: $F_{1,14}$=0.2, p=0.6956 | OFF ChR2 vs. ON ChR2, p=0.0204<br>OFF ChR2 vs. OFF ChR2, p<0.0001<br>OFF YFP vs. ON YFP, p=0.0117<br>OFF YFP vs. OFF YFP, p<0.0001<br>ON ChR2 vs. OFF ChR2, p=0.0004<br>ON YFP vs. OFF YFP, p=0.0048 |
| Figure 2 – figure supplement 1 E | D1-VTA-ChR2=12<br>D1-VTA-YFP=7 | Two-way ANOVA | Stim: $F_{2,22}$=2.8, p=0.0825<br>Group: $F_{1,11}$=0.1, p=0.7707 | |
| Figure 2 – figure supplement 1 F | D1-VTA-ChR2=15<br>D1-VTA-YFP=7 | Two-way ANOVA | Stim: $F_{2,28}$=5.5, p=0.0094<br>Group: $F_{1,14}$=0.3, p=0.6173 | |
| Figure 2 – figure supplement 1 G | D1-VTA-ChR2=15<br>D1-VTA-YFP=7 | Two-way ANOVA | Stim: $F_{2,28}$=4.2, p=0.2<br>Group: $F_{1,14}$=0.2, p=0.6986 | |

| | | | | |
|---|---|---|---|---|
| Figure 2 – figure supplement 1 H | D1-VTA-ChR2=15<br>D1-VTA-YFP=7 | Two-way ANOVA | Stim: $F_{2,28}$=15.1, p<0.001<br>Group: $F_{1,14}$=3.6, p=0.0778 | OFF ChR2 vs. OFF(2) ChR2, p=0.0005<br>OFF ChR2 vs. OFF(2) YFP, p=0.003<br>OFF YFP vs. OFF(2) YFP, p=0.0326 |
| Figure 3 – figure supplement 1 A | D2-VP-ChR2=16<br>D2-VP-YFP=11 | Two-way ANOVA | Stim: $F_{2,36}$=39.1, p<0.001<br>Group: $F_{1,18}$=1.7, p=0.2079 | OFF ChR2 vs. ON ChR2, p<0.0001<br>OFF ChR2 vs. OFF ChR2, p<0.0001<br>OFF ChR2 vs. OFF YFP, p<0.0001<br>OFF YFP vs. ON ChR2, p=0.0006<br>OFF YFP vs. OFF(2) ChR2, p=0.0002<br>OFF YFP vs. OFF(2) YFP, p=0.0020<br>ON ChR2 vs. ON YFP, p=0.0412<br>ON YFP vs. OFF(2) ChR2, p=0.0076 |
| Figure 3 – figure supplement 1 B | D2-VP-ChR2=16<br>D2-VP-YFP=11 | Two-way ANOVA | Stim: $F_{2,34}$=30.5, p<0.001<br>Group: $F_{1,17}$=1.8, p=0.2028 | OFF ChR2 vs. ON ChR2, p<0.0001<br>OFF ChR2 vs. ON YFP, p=0.0390<br>OFF ChR2 vs. OFF(2) ChR2, p<0.0001<br>OFF ChR2 vs. OFF(2) YFP, p<0.0001<br>OFF YFP vs. ON ChR2, p=0.0004<br>OFF YFP vs. OFF(2) ChR2, p=0.0004<br>OFF YFP vs. OFF(2) YFP, p=0.0314<br>ON ChR2 vs. ON YFP, p=0.05<br>ONYFP vs. OFF(2) ChR2, p=0.0325 |
| Figure 3 – figure supplement 1 C | D2-VP-ChR2=18<br>D2-VP-YFP=14 | Two-way ANOVA | Stim: $F_{2,36}$=38.6, p<0.001<br>Group: $F_{1,18}$=0.2, p=0.6564 | OFF ChR2 vs. ON ChR2, p<0.0001<br>OFF ChR2 vs. OFF(2) ChR2, p<0.0001<br>OFF ChR2 vs. OFF(2) YFP, p=0.0036<br>OFF YFP vs. ON ChR2, p=0.0035<br>OFF YFP vs. ON YFP, p=0.0121<br>OFF YFP vs. OFF(2) ChR2, p=0.0173<br>OFF YFP vs. OFF(2) YFP, p<0.0001 |
| Figure 3 – figure supplement 1 D | D2-VP-ChR2=19<br>D2-VP-YFP=14 | Two-way ANOVA | Stim: $F_{2,36}$=33.2, p<0.001<br>Group: $F_{1,18}$=0.02, p=0.8930 | OFF ChR2 vs. ON ChR2, p<0.0001<br>OFF ChR2 vs. OFF(2) ChR2, p<0.0001<br>OFF ChR2 vs. OFF(2) YFP, p=0.0049<br>OFF YFP vs. OFF(2) YFP, p<0.0001 |
| Figure 3 – figure supplement 1 E | D2-VP-ChR2=19<br>D2-VP-YFP=13 | Two-way ANOVA | Stim: $F_{2,36}$=5.8, p=0.0067<br>Group: $F_{1,18}$=1.5, p=0.2387 | OFF ChR2 vs. OFF(2) ChR2, p=0.0419 |
| Figure 3 – figure supplement 1 F | D2-VP-ChR2=18<br>D2-VP-YFP=13 | Two-way ANOVA | Stim: $F_{2,34}$=12.5, p<0.0001<br>Group: $F_{1,17}$=0.1, p=0.7220 | OFF ChR2 vs. OFF(2) ChR2, p=0.0024<br>OFF YFP vs. OFF(2) YFP, p=0.0333 |
| Figure 3 – figure supplement 1 G | D2-VP-ChR2=19<br>D2-VP-YFP=14 | Two-way ANOVA | Stim: $F_{2,36}$=2.0, p=0.1562<br>Group: $F_{1,18}$=0.0, p=0.9379 | |
| Figure 3 – figure supplement 1 H | D2-VP-ChR2=19<br>D2-VP-YFP=14 | Two-way ANOVA | Stim: $F_{2,36}$=33.2, p<0.0001<br>Group: $F_{1,18}$=0.0, p=0.8930 | OFF ChR2 vs. ON ChR2, p<0.0001<br>OFF ChR2 vs. OFF(2) ChR2, p<0.0001<br>OFF ChR2 vs. OFF(2) YFP, p=0.0049<br>OFF YFP vs. OFF(2) YFP, p<0.0001 |
| Figure 4 – figure supplement 1 A | Increase: 13 neurons<br>No Change: 10neurons | One way ANOVA | Increase: $F_{2,24}$=54.1, p<0.0001 | Before vs during, p<0.0001<br>During vs after, p<0.0001 |
| Figure 4 – figure supplement 1 B | Increase: 11 neurons<br>No Change: 2neurons | One way ANOVA | Increase: $F_{2,20}$=19.6, p<0.0001 | Before vs during, p<0.0001<br>During vs after, p=0.0016 |
| Figure 7 – figure supplement 1 A | D2-VP-ChR2-VEH=10<br>D2-VP-ChR2-DIA=110<br>D2-VP-YFP VEH=10<br>D2-VP-YFP DIA=9 | Two-way ANOVA | Stim: F2,18=24.1, p<0.0001<br>Group: $F_{3,27}$=2.7, p=0.0682 | OFF VEH vs. ON ChR2 VEH, p=0.0008<br>OFF ChR2 VEH vs. OFF(2) ChR2 VEH, p=0.0083<br>OFF ChR2 VEH vs. OFF(2) YFP VEH, p=0.0277<br>OFF ChR2 VEH vs. OFF(2) ChR2 VEH, p=0.0087<br>OFF ChR2 VEH vs. OFF(2) YFP VEH, p=0.0277<br>OFF ChR2 VEH vs. OFF:D2-VP-YFP DIA, p=0.0068<br>OFF ChR2 DIA vs. ON ChR2 VEH, p=0.0079<br>OFF ChR2 DIA vs. OFF(2) YFP VEH, p=0.0250<br>OFF ChR2 DIA vs. OFF(2) ChR2 DIA, p=0.0247<br>OFF ChR2 DIA vs. OFF(2) YFP DIA, p=0.0061 |

| | | | | |
|---|---|---|---|---|
| | | | | OFF YFP DIA vs. OFF(2) YFP DIA, p=0.0261<br>ON ChR2 VEH vs. ON ChR2 DIA, p=0.0004<br>ON ChR2 DIA vs. OFF(2) ChR2 VEH, p=0.0035<br>ON ChR2 DIA vs. OFF(2) YFP VEH, p=0.0016<br>ON ChR2 DIA vs. OFF(2) ChR2 DIA, p=0.001<br>ON ChR2 DIA vs. OFF(2) YFP DIA, p=0.0003 |
| Figure 7 – figure supplement 1 B | D2-VP-ChR2-VEH=10<br>D2-VP-ChR2-DIA=10<br>D2-VP-YFP VEH=10<br>D2-VP-YFP DIA=9 | Two-way ANOVA | Stim: F2,18=55.8, p<0.0001<br>Group: $F_{3,27}$=11.6, p<0.0001 | OFF ChR2 VEH vs. ON ChR2 VEH, p<0.0001<br>OFF ChR2 VEH vs. ON YFP DIA, p=0.0488<br>OFF ChR2 VEH vs. OFF(2) ChR2 VEH, p<0.0001<br>OFF ChR2 VEH vs. OFF(2) YFP VEH, p=0.0005<br>OFF ChR2 VEH vs. OFF(2) YFP DIA, p<0.0001<br>OFF YFP VEH vs. ON ChR2 VEH, p<0.0001<br>OFF YFP VEH vs. OFF(2) ChR2 VEH, p<0.0001<br>OFF YFP VEH vs. OFF(2) YFP VEH, p=0.0003<br>OFF VP-YFP VEH vs. OFF(2) YFP DIA, p<0.0001<br>OFF YFP DIA vs. ON ChR2 VEH, p<0.0001<br>OFF YFP DIA vs. ON YFP DIA, p=0.0294<br>OFF YFP DIA vs. OFF(2) ChR2 VEH, p<0.0001<br>OFF YFP DIA vs. OFF(2) YFP VEH, p=0.0005<br>OFF YFP DIA vs. OFF(2) YFP DIA, p<0.0001<br>OFF ChR2 DIA vs. ON ChR2 VEH, p<0.0001<br>OFF ChR2 DIA vs. ON YFP DIA, p=0.0488<br>OFF ChR2 DIA vs. OFF(2) ChR2 VEH, p<0.0001<br>OFF ChR2 DIA vs. OFF(2) YFP VEH, p=0.0005<br>OFF ChR2 DIA vs. OFF(2) YFP DIA, p<0.0001<br>ON ChR2 VEH vs. ON YFP VEH, p=0.0197<br>ON ChR2 VEH vs. ON ChR2 DIA, p<0.0001<br>ON ChR2 VEH vs. OFF(2) ChR2 DIA, p=0.0006<br>ON ChR2 DIA vs. OFF(2) ChR2 VEH, p<0.0001<br>ON ChR2 DIA vs. OFF(2) YFP VEH, p=0.0045<br>ON ChR2 DIA vs. OFF(2) YFP DIA, p=0.0003<br>OFF ChR2 VEH vs. OFF(2) ChR2 DIA, 0.0022<br>OFF YFP DIA vs. OFF(2) ChR2 DIA, p=0.041 |
| Figure 7 – figure supplement 1 C | D2-VP-ChR2-VEH=8<br>D2-VP-ChR2-DIA=10<br>D2-VP-YFP VEH=8<br>D2-VP-YFP DIA=9 | Two-way ANOVA | Stim: F2,18=80.7, p<0.0001<br>Group: $F_{3,27}$=0.3, p=0.8511 | OFF ChR2 VEH vs. ON ChR2 VEH, p<0.0001<br>OFF ChR2 VEH vs. ON ChR2 DIA, p=0.0005<br>OFF ChR2 VEH vs. OFF(2) ChR2 VEH, p=0.0023<br>OFF ChR2 VEH vs. OFF YFP VEH, p=0.0024<br>OFF ChR2 VEH vs. OFF(2) YFP DIA, p=0.0008<br>OFF YFP VEH vs. ON ChR2 VEH, p<0.0001<br>OFF YFP VEH vs. ON YFP VEH, p=0.001<br>OFF YFP VEH vs. ON ChR2 DIA, p<0.0001<br>OFF YFP VEH vs. OFF(2) ChR2 VEH, p=0.0077<br>OFF YFP VEH vs. OFF(2) YFP VEH, p<0.0001<br>OFF YFP VEH vs. OFF(2) ChR2 DIA, p=0.0068<br>OFF YFP VEH vs. OFF(2) YFP DIA, p<0.0001<br>OFF ChR2 DIA vs. ON ChR2 VEH, p<0.0001<br>OFF ChR2 DIA vs. ON YFP VEH, p=0.0049<br>OFF ChR2 DIA vs. ON ChR2 DIA, p<0.0001<br>OFF ChR2 DIA vs. ON YFP DIA, p=0.0042<br>OFF ChR2 DIA vs. OFF(2) ChR2 VEH, p=0.0005<br>OFF ChR2 DIA vs. OFF(2) YFP VEH, p<0.0001<br>OFF ChR2 DIA vs. OFF(2) ChR2 DIA, p<0.0001<br>OFF ChR2 DIA vs. OFF(2) YFP DIA, p<0.0001<br>OFF YFP DIA vs. ON ChR2 VEH, p=0.0001<br>OFF YFP DIA vs. ON ChR2 DIA, p=0.0001<br>OFF YFP DIA vs. ON YFP DIA, p=0.0092<br>OFF YFP DIA vs. OFF(2) ChR2 VEH, p=0.0388<br>OFF YFP DIA vs. OFF(2) YFP VEH, p=0.0008<br>OFF YFP DIA vs. OFF(2) ChR2 DIA, p=0.0358<br>OFF YFP DIA vs. OFF(2) YFP DIA, p<0.0001 |
| Figure 7 – figure supplement 1 D | D2-VP-ChR2-VEH=8<br>D2-VP-ChR2-DIA=10<br>D2-VP-YFP VEH=8<br>D2-VP-YFP DIA=9 | Two-way ANOVA | Stim: F2,16=132.2, p<0.0001<br>Group: $F_{3,24}$=0.7, p=0.5443 | OFF ChR2 VEH vs. ON ChR2 VEH, p<0.0001<br>OFF ChR2 VEH vs. ON YFP VEH, p=0.0182<br>OFF ChR2 VEH vs. ON ChR2 DIA, p<0.0001<br>OFF ChR2 VEH vs. ON YFP DIA, p=0.0031<br>OFF ChR2 VEH vs. OFF(2) ChR2 VEH, p<0.0001 |

| | | | | |
|---|---|---|---|---|
| | | | | OFF ChR2 VEH vs. OFF(2) YFP VEH, p=0.0002<br>OFF ChR2 VEH vs. OFF(2) ChR2 DIA, p=0.0287<br>OFF ChR2 VEH vs. OFF(2) YFP DIA, p=0.0008<br>OFF YFP VEH vs. ON:D2-VP-ChR2 VEH, p<0.0001<br>OFF YFP VEH vs. ON YFP VEH, p<0.0001<br>OFF YFP VEH vs. ON ChR2 DIA, p<0.0001<br>OFF YFP VEH vs. ON YFP DIA, p<0.0001<br>OFF YFP VEH vs. OFF(2) ChR2 VEH, p<0.0001<br>OFF YFP VEH vs. OFF(2) YFP VEH, p<0.0001<br>OFF YFP VEH vs. OFF(2) ChR2 DIA, p<0.0001<br>OFF YFP VEH vs. OFF(2) YFP DIA, p<0.0001<br>OFF ChR2 DIA vs. ON ChR2 VEH, p<0.0001<br>OFF ChR2 DIA vs. ON YFP VEH, p=0.0017<br>OFF ChR2 DIA vs. ON ChR2 DIA, p<0.0001<br>OFF ChR2 DIA vs. ON YFP DIA, p=0.0002<br>OFF ChR2 DIA vs. OFF(2) ChR2 VEH, p=0.0002<br>OFF ChR2 DIA vs. OFF(2) YFP VEH, p<0.0001<br>OFF ChR2 DIA vs. OFF(2) ChR2 DIA, p<0.0001<br>OFF ChR2 DIA vs. OFF(2) YFP DIA, p<0.0001<br>OFF YFP DIA vs. ON ChR2 VEH, p<0.0001<br>OFF YFP DIA vs. ON YFP VEH, p=0.0158<br>OFF YFP DIA vs. ON ChR2 DIA, p<0.0001<br>OFF YFP DIA vs. ON YFP DIA, p<0.0001<br>OFF YFP DIA vs. OFF(2) ChR2 VEH, p=0.002<br>OFF YFP DIA vs. OFF(2)YFP VEH, p=0.0001<br>OFF YFP DIA vs. OFF(2) ChR2 DIA, p=0.0253<br>OFF YFP DIA vs. OFF(2) YFP DIA, p<0.0001 |
| Figure 7 – figure supplement 1 E | D2-VP-ChR2-VEH=9<br>D2-VP-ChR2-DIA=8<br>D2-VP-YFP VEH=8<br>D2-VP-YFP DIA=9 | Two-way ANOVA | Stim: F2,16=0.1, p=9054<br>Group: $F_{3,24}$=0.04, p=0.9871 | |
| Figure 7 – figure supplement 1 F | D2-VP-ChR2-VEH=9<br>D2-VP-ChR2-DIA=9<br>D2-VP-YFP VEH=8<br>D2-VP-YFP DIA=9 | Two-way ANOVA | Stim: F2,16=28.2, p<0.0001<br>Group: $F_{3,24}$=0.09, p=0.9668 | OFF CHR2 DIA vs. ON CHR2 DIA, p=0.0436<br>OFF CHR2 DIA vs. OFF(2) CHR2 DIA, p=0.011<br>OFF CHR2 VEH vs. OFF(2) CHR2 VEH, p=0.0348 |
| Figure 7 – figure supplement 1 G | D2-VP-ChR2-VEH=9<br>D2-VP-ChR2-DIA=9<br>D2-VP-YFP VEH=8<br>D2-VP-YFP DIA=9 | Two-way ANOVA | Stim: F2,16=286.8, p<0.0001<br>Group: $F_{3,24}$=0.04, p=0.9870 | OFF YFP VEH vs. OFF YFP VEH, p=0.0393<br>OFF CHR2 DIA vs. OFF(2) CHR2 DIA, p=0.0033<br>OFF CHR2 VEH vs. OFF(2) CHR2 VEH, 0.0438 |
| Figure 7 – figure supplement 1 H | D2-VP-ChR2-VEH=9<br>D2-VP-ChR2-DIA=9<br>D2-VP-YFP VEH=9<br>D2-VP-YFP DIA=9 | Two-way ANOVA | Stim: F2,16=151.0, p<0.0001<br>Group: $F_{3,24}$=0.2, p=0.8636 | OFF YFP VEH vs. OFF(2) YFP VEH, p=0.0005<br>OFF YFP VEH vs. OFF(2) CHR2 DIA, p=0.0002<br>OFF YFP DIA vs. OFF(2) YFP VEH, p=0.0022<br>OFF YFP DIA vs. OFF(2) YFP DIA, p=0.0028<br>OFF YFP DIA vs. OFF(2) CHR2 DIA, p=0.0052<br>OFF YFP DIA vs. OFF(2) CHR2 VEH, p=0.0168<br>OFF CHR2 DIA vs. OFF(2) YFP VEH, p<0.0001<br>OFF CHR2 DIA vs. OFF(2) YFP DIA, p=0.018<br>OFF CHR2 DIA vs. OFF(2) CHR2 DIA, p<0.0001<br>OFF CHR2 DIA vs. OFF(2) CHR2 VEH, p=0.0109<br>OFF CHR2 VEH vs. OFF(2) YFP VEH, p=0.0074<br>OFF CHR2 VEH vs. OFF(2) YFP DIA, p=0.0102<br>OFF CHR2 VEH vs. OFF(2) CHR2 DIA, p=0.0006<br>OFF CHR2 VEH vs. OFF(2) CHR2 VEH, p=0.0007 |
